# Characterizing Photosynthetic Biofuel Production: Isotopically non-stationary ^13^C metabolic flux analysis (INST-^13^CMFA) on limonene producing *Synechococcus* sp. PCC 7002

**DOI:** 10.1101/2022.03.30.486112

**Authors:** Darrian M. Newman, Cara L. Sake, Alex J. Metcalf, Fiona K. Davies, Nanette R. Boyle

**Affiliations:** Department of Chemical and Biological Engineering, Colorado School of Mines, Golden, CO 80401, USA; Living Ink Technologies, Commerce City, CO 80022, USA

**Author notes:** These authors have contributed equally to this work and share first authorship (Darrian M. Newman, Cara L. Sake). **Correspondence:** Nanette Boyle.

**Keywords:** cyanobacteria, terpenoid, phosphoenolpyruvate carboxylase, malic enzyme, ATP:NADPH ratio, reductive TCA cycle

## Abstract

*Synechococcus* sp. PCC 7002 is a unicellular cyanobacterium capable of fast growth, even under high light intensity and high salinity. These attributes along with genetic tractability make *Synechococcus* sp. PCC 7002 an attractive candidate for industrial scale production of specialty and commodity chemicals. One such strain produces limonene, an energy dense diesel jet fuel drop-in additive, at a titer of 4 mg/L over a four-day incubation period. In this study, we use the state-of-the-art whole-cell characterization tool, isotopically non-stationary ^13^C metabolic flux analysis (INST-^13^CMFA) to determine intracellular fluxes through the pathways of central metabolism for the limonene producing strain and wild type strain of *Synechococcus* sp. PCC 7002. We find similar flux distribution in the Calvin-Benson-Bassham cycle, photorespiration, oxidative pentose phosphate pathway, and reductive tricarboxylic acid cycle. The key difference between strains is observed in the production of pyruvate. The limonene producing strain displays significantly higher flux through the amphibolic pathways of phosphoenolpyruvate carboxylase and the malic enzyme to synthesize pyruvate, while the wild type strain uses pyruvate kinase in a single step. Our findings suggest that this flux distribution is a mechanism to recover a physiologically optimal ratio of ATP to NADPH. The upregulation of this amphibolic pathway may act to restore the physiological ATP:NADPH ratio that has been disturbed by limonene biosynthesis.

## 1 Introduction

Amid growing concerns over climate change and increasing anthropogenic carbon emissions, cyanobacteria have emerged as a promising platform for the sustainable production of a wide range of specialty and commodity chemicals. Cyanobacteria utilize the Calvin-Benson-Bassham (CBB) cycle to fix atmospheric carbon dioxide into biomass, metabolites and important industrially relevant chemicals and fuels using only light and trace minerals as additional inputs. Cyanobacterial cell factories are naturally capable of producing a wide variety of products: sugars, alcohols, acids, alkanes, alkenes, ketones, fatty acids, and terpenoids (Khan et al., 2019). The efficacy of sustainable chemical production by cyanobacteria is strengthened by their genetic tractability as well as higher photosynthetic efficiency and growth rates compared to land plants (Ducat et al., 2011;Berla et al., 2013). *Synechococcus* sp. PCC 7002 (hereafter *Synechococcus* 7002) is an attractive candidate for production of sustainable industrially relevant chemicals to displace petrochemicals due to a short doubling time of only 2.6 h when provided reduced nitrogen (Ludwig and Bryant, 2012). *Synechococcus* 7002 can also tolerate high salinity and light intensity (Batterton and Baalen, 1971;Nomura et al., 2006).

Two strains are studied here, the wild type (WT) *Synechococcus* 7002 strain and a limonene producing (LS) strain, engineered by Davies et al. (Davies et al., 2014). The *Synechococcus* 7002 LS strain produces 4 mg L^-1^ of the terpenoid limonene over a 96 hour growth period (Davies et al., 2014). Briefly, L-limonene is synthesized in cyanobacterial cells via the methylerythritol 4-phosphate (MEP) pathway, a linear seven step pathway beginning with the condensation of glyceraldehyde 3-phosphate (GAP) and pyruvate (PYR), and ending with the production of either isopentenyl diphosphate (IPP) or dimethylallyl diphosphate (DMAPP) along with the expression of a heterologous enzyme (limonene synthase). Limonene is an energy dense molecule, which is why it is an attractive candidate as a drop-in biodiesel jet fuel additive. Few studies have been conducted to understand the energetic burden on cyanobacterial metabolism when producing such energy dense molecules as limonene. Many attempts, however, have been made to increase flux through the MEP pathway toward terpenoids such as limonene through the overexpression of bottlenecks (Gao et al., 2016;Englund et al., 2018) and by alleviating competition for carbon with sinks such as glycogen (Davies et al., 2014;Hendry et al., 2017). However, titers of terpenoid products including limonene by *Synechococcus* 7002 LS remain too low to be industrially relevant and economical producers of biofuel molecules.

The allocation of carbon through cyanobacterial metabolism and consequences of the additional energetic burden must be understood if we are to significantly increase production of terpenoids. Isotopically nonstationary ^13^C metabolic flux analysis (INST-^13^CMFA) is a state-of-the-art tool that we utilized to investigate these questions. INST-^13^CMFA uses isotopically labeled carbon to produce time-dependent mass isotopomer distributions (MIDs) of key metabolites in central metabolism, from which flux values can be estimated for an entire network of reactions by iteratively solving the flux network and comparing to the experimental data provided. INST-^13^CMFA has been utilized to elucidate metabolic phenotypes in *Synechococcus* 7002 (Hendry et al., 2017;Qian et al., 2018;Abernathy et al., 2019), *Synechocystis* sp. PCC 6803 (You et al., 2014;Adebiyi et al., 2015;Nakajima et al., 2017;Yu King Hing et al., 2019), and *Synechococcus Elongatus* PCC 7942 (or closely related species UTEX 2973) (Abernathy et al., 2017;Jazmin et al., 2017;Cheah et al., 2020). In this study, we use INST-^13^CMFA to characterize the central metabolism of the *Synechococcus* 7002 WT and LS strains to glean information about phenotypic differences induced by the production of limonene. We find significant carbon redistribution in central carbon metabolism between the two strains. This redistribution of carbon flux appears to be induced not by the need for carbon reallocation, but for recovering the physiologically optimal ratio of ATP to NADPH needed for biomass accumulation.

## 2 Methods

### 2.1 Strains and cultivation conditions

*Synechococcus* 7002 cultures were obtained from the American Type Culture Collection and the limonene producing strain (LS) was provided by Dr. Davies (Davies et al., 2014). *Synechococcus* 7002 cultures were grown in BG11 medium supplemented with 8.26 mM Trizma base (chemicals supplied by Sigma-Aldrich, unless otherwise noted) titrated to a pH of 8.2 and maintained on solid BG11 medium plates containing 1.5% (w/v) agar (Difco) plates. A final concentration of 50 μg/mL spectinomycin was added to cultures of the limonene producing strain. Liquid cultures were propagated and grown in 250 mL and 500 mL beveled Erlenmeyer flasks with 100 mL and 200 mL of media, respectively. Flasks were shaken at 180 rpm in an INFORS HT Minitron shake plate incubator at ambient CO_2_ conditions (<0.1%), and 37 °C. The approximate light intensity was 80 μmol photons m^-2^ s^-1^ in continuous illumination by white fluorescent bulbs. Cell growth was monitored using spectrophotometric optical density (OD_730_).

### 2.2 Dynamic labeling and quenching experiment

The labeling experiment was performed on a benchtop flask shaker under 20 μmol photon m^-2^ s^-1^ light from white fluorescent lights. The cultures were first grown up to mid-exponential phase (OD_730_~0.8) at the conditions previously described. Before the addition of label, 15 mL of culture was pipetted out and immediately quenched in 30 mL of partially frozen saline quench solution in a 50 mL conical centrifuge tube, as described in Sake et al.(Sake et al., 2020). This sample corresponded to the zero timepoint sample (t=0). Then a 5 mL bolus of 0.8 M ^13^C sodium bicarbonate (98 atom % ^13^C, 99% purity, Sigma-Aldrich, USA) was injected into the flask at time t=0. 15 mL Samples were rapidly quenched at 20s, 40s, 1 min, 2min, 4min, 6min, 10min, 15min, 20min, 40min, 60min. The quenched samples were centrifuged in an Eppendorf 5810R swinging bucket centrifuge at 4,000 rpm for 15 minutes at −2°C, and the supernatant was decanted. The pellet was washed in 2 mL chilled quenching solution, transferred to a 2 mL microcentrifuge tube, and centrifuged again at 8,000*xg* on a tabletop centrifuge at −2°C. The pellet was stored at −20°C for later extraction.

### 2.3 Metabolite extraction

Intracellular metabolites were extracted from the frozen cell pellets using a methanol extraction (Sake et al., 2020). Cell pellets were resuspended in 500 μL pure methanol and spiked with ribitol and PIPES internal standards for final concentrations of 150 ppb and 50 ppb, respectively. Samples were frozen in liquid nitrogen, thawed on ice, and vortexed at 0°C for 5 minutes and 1000 rpm. This freeze/thaw/vortex cycle was repeated twice more, and samples were centrifuged at 8000*x*g for 5 minutes at −9°C in a Sorvall Legend Micro 17R (Thermo Scientific). The supernatant (extract) was collected in a new tube and stored at −20°C. The extraction process was repeated twice more with 500 μL of 50% methanol, each time adding the collected extract to the first portion. All samples were then dried overnight under vacuum at 45°C in a Savant SPD131DDA SpeedVac (Thermo Scientific). Dried extracts were resuspended in 150 μL optima water and filtered with nylon filter tubes (Spin-X, Costar). Filters were rinsed with an additional 50 μL optima water for a total concentrated extract volume of 200 μL. Finally, the samples were filtered through a 3 kilodalton filter (Spin Filter 3K, VWR) and collected for LC-MS/MS analysis.

### 2.4 LC/MS-MS quantitation of metabolites

Metabolite extracts were analyzed using an LC-MS/MS method adapted from Young et al. (Young et al., 2011). Analysis was performed using a Phenomenex 150 mm x 2 mm Synergi Hydro-RP column connected to an Agilent 1200 Series HPLC system and autosampler in tandem with the AB Sciex 5500 QTrap MS/MS system. LC was performed with an injection volume of 20 μL, using gradient elution of 10 mM tributylamine and 15 mM acetic acid (aqueous phase) with acetonitrile (organic phase) at a constant flow of 0.3 mL/min. The gradient profile of the organic phase is as follows: 0% B (0 min), 8% B (8 min), 16% B (15 min), 30% B (16.5 min), 30% B (19 min), 90% B (21.5 min), 90% B (26.5 min), 0% B (26.6 min), and 0% B (30.5 min). MS analysis was performed in negative ionization mode using a multiple reaction monitoring (MRM) acquisition method. Data acquisition was performed on the Sciex Analyst 1.7 software. Metabolite pool sizes were quantified using Sciex MultiQuant 3.0.3 software. MSConvert was used to process data files into an open-source format, and isotope labeling profiles were processed using a combination of pyOpenMS and SciPy packages in Python.

### 2.5 Isotopically non-stationary ^13^C metabolic flux analysis

We used a previously published model of the central metabolic network of *Synechococcus* 7002 (Adebiyi et al., 2015;Abernathy et al., 2017;Hendry et al., 2017;Abernathy et al., 2019). Models for the wild type and LS strains were constrained to match the composition reported by Abernathy et al. for *Synechococcus* 7002 in photoautotrophic conditions (Abernathy et al., 2019). We assumed the biomass formation equation to remain constant between strains, given the low limonene titers (4 mg/L) found by Davies et al. (Davies et al., 2014). We constrained each network to the biomass accumulation rate calculated during the growth phase of the experiment (Figure S2).

The flux network and atom transitions for this study were taken from previous studies on *Synechococcus* 7002 and closely related strains (Adebiyi et al., 2015;Abernathy et al., 2017;Hendry et al., 2017;Abernathy et al., 2019), including in the network the Calvin-Benson-Bassham cycle, photorespiration pathway, oxidative pentose phosphate pathway, tricarboxylic acid (TCA) cycle, and amphibolic reactions. The full metabolic network and atom transitions can be found in Table S3. The MEP condensed pathway to limonene was constructed based on gene annotations from the KEGG database (Kanehisa, 2002;Kanehisa et al., 2014). We used the lumped biomass equation constructed Abernathy et al. (Abernathy et al., 2019). from biomass composition analysis. The MATLAB-based INCA toolbox (Young, 2014) was used to construct the network, and INST-MFA simulations were run to estimate reaction fluxes and metabolite pool sizes by minimizing the difference between simulated and measured mass isotopomer distributions provided to the model. Data used to produce MID figures in Figure S1 are tabulated in Table S1 and S2. The parameter continuation method provided by INCA estimated 95% confidence intervals around each estimated parameter. Dilution parameters were applied to metabolites as needed to account for labeling dilutions from metabolically inactive pools (Young et al., 2011) (Table S3 & S4).

## 3 Results and discussion

### 3.1 Differential ^13^C enrichment

Isotopic labeling was measured in 11 key metabolites (see Figure 1), and time dependent mass isotopomer distributions show the dynamics of metabolite labeling (Figure S1). To account for low enrichment, dilution pools were provided to the model for 3PG, Ru5P, PEP, GAP, and DHAP (Table S3). Dilution pools represent mechanisms in the cell that prevent certain metabolite pools from reaching expected enrichment, due to phenomena like metabolite channeling that cause intermediates to proceed through multiple reactions without mixing with entire metabolite pool (Ishikawa et al., 2004;Broddrick et al., 2016;Abernathy et al., 2019).

**Figure 1.**
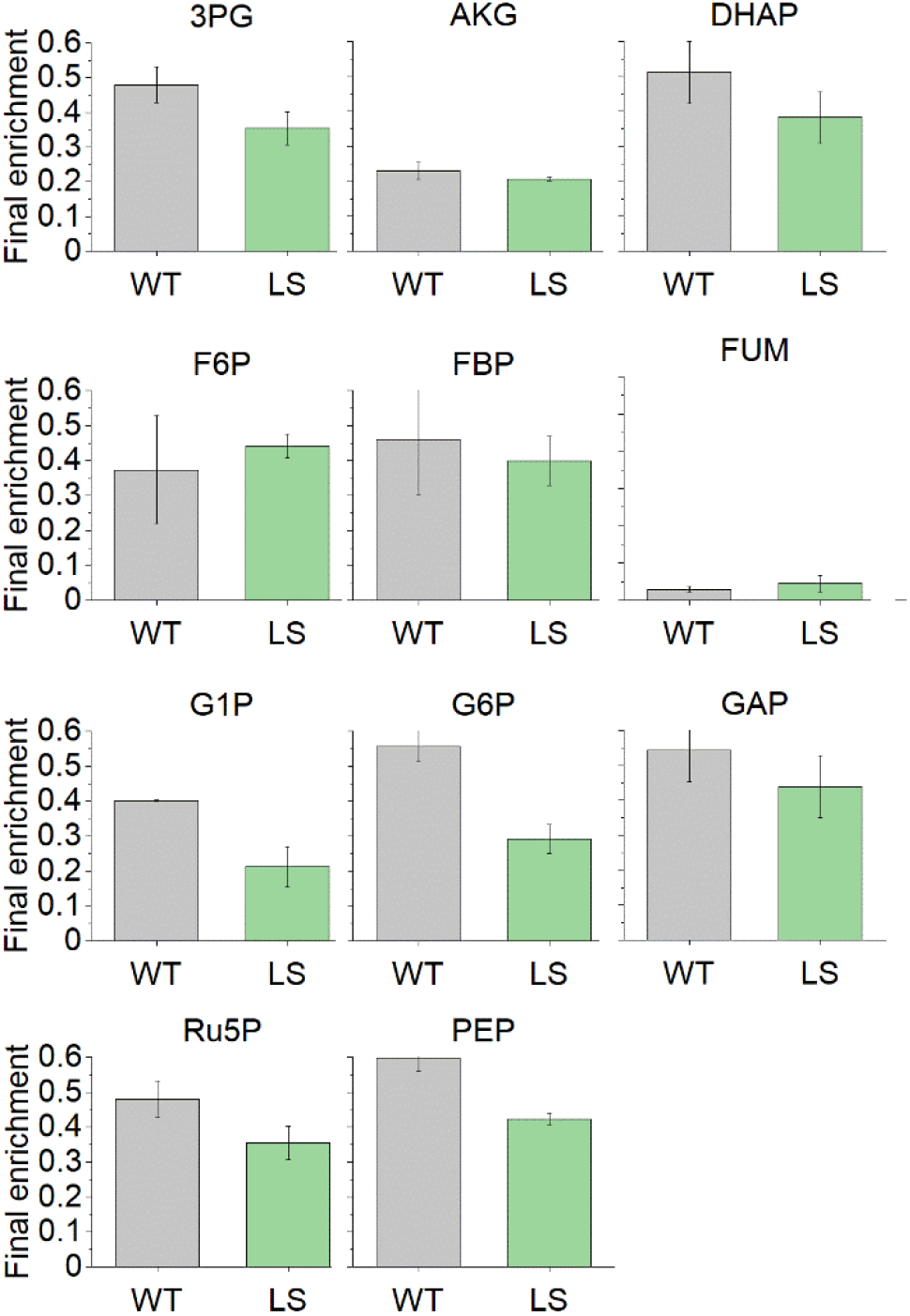
Final ^13^C enrichment of the 11 metabolites used to inform the flux distributions through the *Synechococcus* 7002 central metabolic network. Final enrichment values correspond to MID data at the 60 minute timepoint (Figure S1). Error bars represent standard error (n=3). The final enrichment of PEP was found to be significantly different between the WT and LS strains (p=0.01144).

We noticed interesting behavior in the PEP node. The final enrichment for each strain (Figure 1) shows a lower overall enrichment of the PEP metabolite pool in the LS strain compared to the WT strain. In the WT strain we measured the steady state PEP enrichment of nearly 60%, significantly different from the LS enrichment of 42% (p=.01144). This difference necessitated the use of a dilution pool for the LS PEP pool in the INCA model to account for the low labeling and was needed for the SSR of the solved flux map to fall within the accepted error range.

### 3.2 Flux measurements for the wild type and limonene producing strain

The WT flux map was solved by the INCA package in MATLAB to an acceptable sum of squared residuals (SSR) of 482.1, within the range of 472.8 to 601.0. Likewise, the LS flux map was solved to an acceptable SSR of 538.4 within the range of 435.2 to 558.5. The resulting flux maps were assigned 95% confidence intervals for each individual reaction in the network (Table S4). The general distribution of flux throughout central metabolism concurs with previous INST-MFA studies on *Synechococcus* 7002 and similar cyanobacterial strains (Adebiyi et al., 2015;Abernathy et al., 2017;Hendry et al., 2017;Abernathy et al., 2019), with a large proportion of flux directed through the CBB cycle for carbon fixation, relatively low flux through the TCA cycle, and similar carbon allocation toward glycogen for storage (Figure 2).

**Figure 2.**
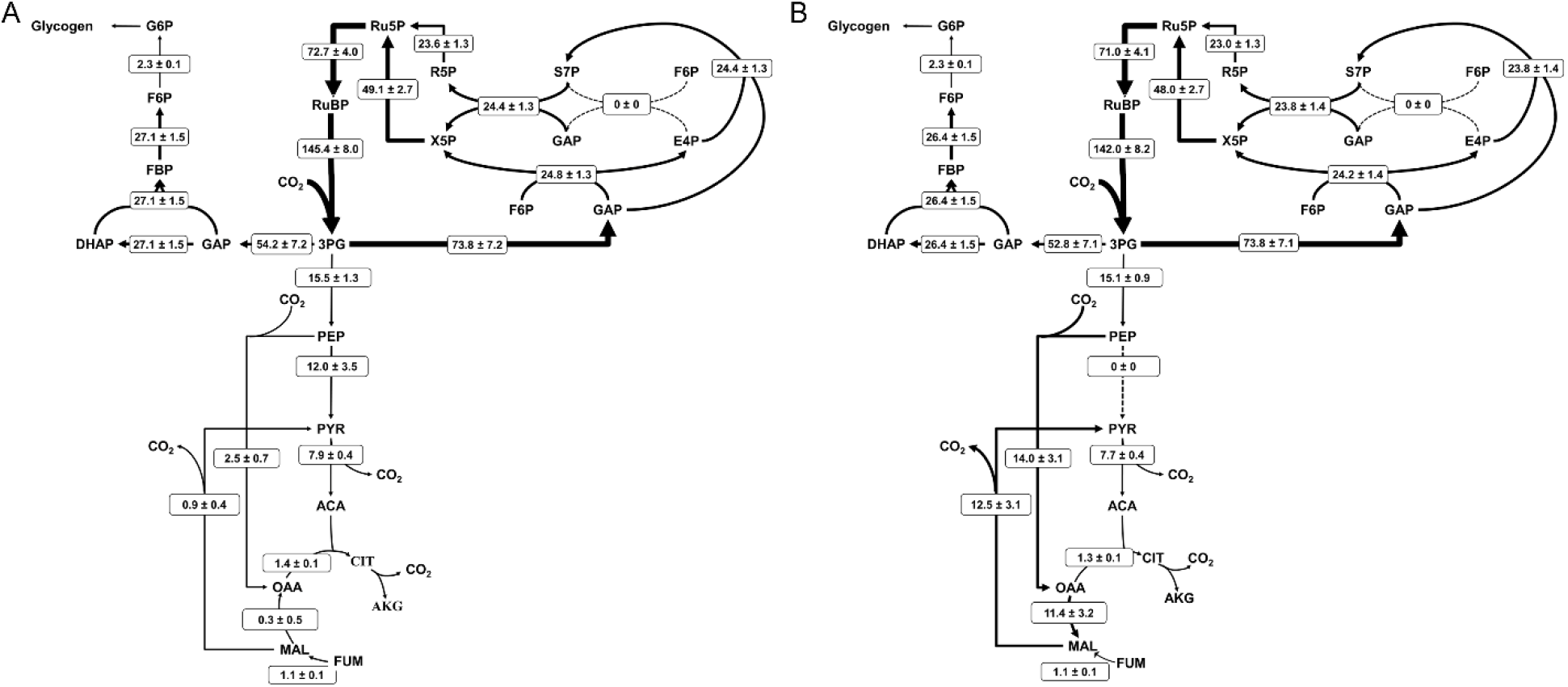
*Metabolic* flux maps for *Synechococcus* 7002 (A) WT and (B) LS. Net fluxes are shown in the form M ± SE where M is the calculated flux and SE is the standard error of the 95% confidence interval, calculated using upper and lower flux bounds from parameter continuation analysis. Arrow thickness is proportional to flux through the reaction, and dotted lines indicate no significant flux. Flux values for every reaction in the network, upper and lower bounds of their respective 95% confidence intervals, and dilution parameters are listed in Table S4.

Our model allowed for an active photorespiration pathway, but the flux was determined to be negligible. This is likely due to the high bicarbonate concentration accumulated inside the cells after a bolus of ^13^C bicarbonate was added at the zero timepoint. Cyanobacteria are able to rapidly transport available bicarbonate into the cell (Colman, 1989), validating both the suppression of photorespiration as well as the establishment of an intracellular dissolved inorganic carbon pool dominated by ^13^C. Both ^13^C and ^12^C CO_2_ pools were made available to the cell for transport in simulations, but only the ^13^C pool was utilized by the network (Table S2).

### 3.3 Metabolic flux analysis reveals increased flux through phosphoenolpyruvate carboxylase

The solved flux maps describe differing usage of the two separate pyruvate synthesis pathways. The one step pathway from PEP to PYR proceeds through pyruvate kinase (PK), reacting ADP and PEP together to form ATP and PYR. Alternatively, phosphoenolpyruvate carboxylase (PEPc), malate dehydrogenase (MDH), and the malic enzyme (ME), synthesize PYR from PEP in an amphibolic loop by first reacting PEP and CO_2_ to OAA at the expense of an ATP, reducing OAA to MAL at the expense of the oxidation of NADH to NAD^+^, before decarboxylating and oxidizing MAL to PYR while reducing NADP^+^ to NADPH (Figure 3). The three-step pathway forms part of the reductive TCA cycle. In the bifurcated structure of the cyanobacterial TCA cycle, the reductive portion supports higher flux than the oxidative portion, which is suppressed in light conditions (Zhang and Bryant, 2011;Xiong et al., 2017), typically with low flux through TCA intermediates like CIT and AKG, and more flux supplying OAA and MAL directly from PEP (Iijima et al., 2021), additionally supplemented by FUM through purine synthesis (Knoop et al., 2010). As expected, we see low fluxes in Figure 2 for the oxidative TCA cycle, with flux driven by the production of biomass precursors (ie. AKG).

**Figure 3.**
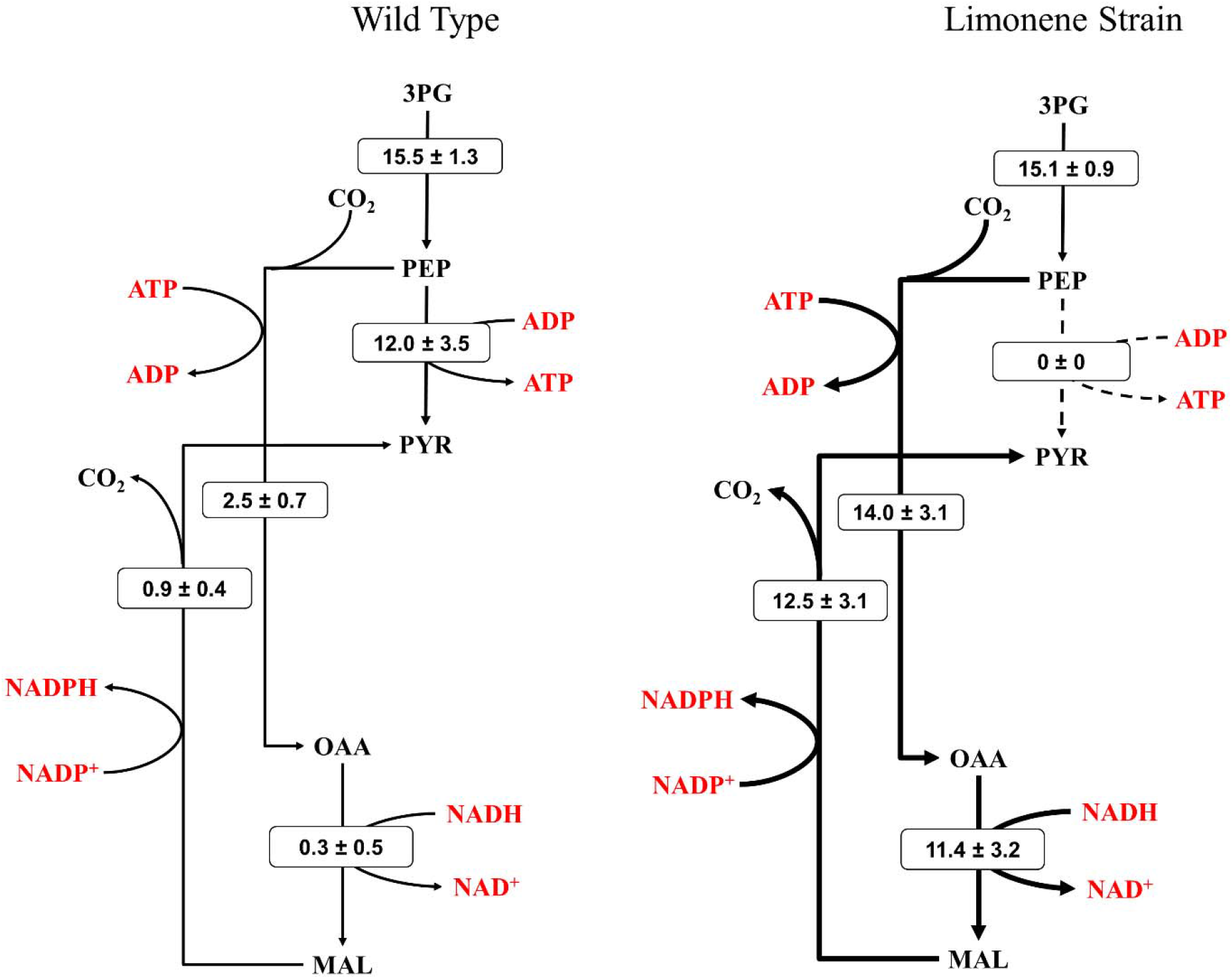
Fluxes of pyruvate synthesis from phosphoenolpyruvate. Flux distribution between the one step, pyruvate kinase mediated reaction from PEP to PYR versus the three step amphibolic pathway, involving phosphoenolpyruvate carboxylase, malate dehydrogenase, and the malic enzyme differ significantly between WT and LS strains. Energy molecule cofactors aiding in each reaction are depicted in red. Thickness of the black arrows are proportional to relative flux values, provided in the form M ± SE where M is the calculated flux and SE is the standard error of the 95% confidence interval.

The flux maps reveal significant differences in the reductive TCA cycle between the WT and LS strains. In the WT strain, PEPc fixes 2.5±0.7 umol/gDW/hr CO_2_ into oxaloacetate, of which only 0.9±0.4 umol/gDW/hr is cycled through the malic enzyme, resulting minimal production of PYR from PEP compared to the PK route. The limonene strain fixes 14.0±3.1 umol/gDW/hr of CO_2_ by PEPc activity, ultimately sending higher flux through the malic enzyme as well, 12.5±3.1 umol/gDW/hr. This significant difference in flux distribution is the direct result of differential labeling at the PEP node, which can likely be attributed to metabolite channeling. Labeling suggests that the PEP pool is more homogenous in the WT strain, where PYR production is dominated by the PK route. Alternatively, metabolite channeling through PEPc and neighboring enzymes in the amphibolic loop prevent PEP directed toward MAL from mixing homogenously within the cell. INST -^13^C-MFA has already been used to identify subcellular organization and provide evidence of metabolite channeling on metabolite labeling patterns in *Synechococcus* 7002 (Abernathy et al., 2019). The traditional view of prokaryotic cyanobacteria as spatially unorganized with no subcellular compartmentalization has been challenged by other studies as well (Ishikawa et al., 2004;Broddrick et al., 2016).

### 3.4 Global regulation of the ATP:NADPH balance may be responsible for flux redistribution

Limonene is an energy dense molecule, with a molecular formula of C_10_H_16_, including two double bonds. To biosynthesize one mole of limonene through the native cyanobacterial MEP pathway, 36 moles of ATP and 28 moles of NADPH are consumed, a 1.29 ATP:NADPH ratio. This ratio of consumption is almost exactly the ratio in which ATP and NADPH are produced from linear electron flow in photosynthetic systems, a 9:7 ratio of ATP to NADPH (Alric et al., 2010;Kramer and Evans, 2011). However, it is estimated that the optimal ATP:NADPH ratio for biomass accumulation in cyanobacteria is greater than 1.5 (Alric et al., 2010;Erdrich et al., 2014;Hendry et al., 2016). Alternative electron flows account for the additional production of energy molecules, and are tightly regulated (Trost and Lemaire, 2013;Wilde and Hihara, 2016).

Limonene synthesis requires a higher input of NADPH compared to ATP, relative to the optimal ratio required for biomass accumulation. We hypothesize that even at small titers (4 mg L^-1^) this disruption of the relative abundance of ATP to NADPH trigger a global regulatory response aimed at recovering the optimal ratio. While PYR synthesis by PK produces an ATP, expending an ATP by rerouting carbon down PEPc results in the production of NADPH, at the additional cost of a molecule of NADH as well, from the reduction of OAA to MAL. This exchange of ATP and NADH for NADPH counteracts the relatively high NADPH consumption needed for limonene synthesis. Although regulation of cyanobacterial NADPH homeostasis is generally poorly understood (Ishikawa and Kawai-Yamada, 2019), the malic enzyme route is known in bacteria and archaea as an NADPH generating reaction (Singh et al., 2008;Negi et al., 2015;Spaans et al., 2015).

Interestingly, in other studies conducted on terpenoid production strains it is observed that the introduction of terpenoid based carbon sinks do not increase photosynthetic efficiency (Wang et al., 2016). This contrasts findings in which sucrose production in cyanobacteria is accompanied by increases in photosynthetic efficiency (Abramson et al., 2016;Abramson et al., 2018;Lin et al., 2020). It is reasonable to assume that the increase in photosynthetic efficiency in response to the addition of a sucrose carbon sink can be attributed to the fact that sucrose production does not affect intracellular ATP:NADPH equilibrium (Thiel et al., 2019). Production of molecules like ethanol, isobutanol, lactate, isoprene, and many others require a much lower ATP:NADPH ratio for biosynthesis than the optimal cellular ratio for biomass accumulation, similar to limonene.

Our findings suggest that limonene is indeed causing a cellular imbalance of ATP:NADPH, and physiological changes are occurring in central metabolism in order to correct this imbalance. We observe the conversion of PEP to PYR as an important control mechanism to tune the ATP:NADPH ratio back to the physiologically optimal value, allowing the LS strain to recover biomass accumulation rates similar to the WT strain. We show that the effects of genetic modifications to the MEP pathway are not confined there, but drive flux redistribution in central metabolic pathways, such as PK, PEPc and the malic enzyme. More research is needed to elucidate the regulatory mechanisms controlling the two pathways for PYR conversion from PEP.

INST-^13^CMFA has also provided direction for future experiments to study the effects of pyruvate synthesis on biofuel production strains. Optimizing the distribution of flux from the PEP node through overexpression or suppression of PEPc and PK could increase limonene titers and lead to increased photosynthetic efficiency through optimization of the ATP:NADPH ratio. Alternatively, co-expression of enzymes for products that require a relatively high ATP:NADPH ratio could offset the disturbance caused by limonene production.

## 4 Conclusions

To characterize the phenotype of a limonene producing strain of *Synechococcus* 7002, INST-^13^C-MFA was performed on the LS strain, and compared to the results of the WT strain. Results depicted similar flux distribution in the CBB cycle, gluconeogenesis, and the oxidative TCA cycle, however flux maps revealed differential flux through pyruvate biosynthesis pathways. The LS strain redirects carbon flux from PEP to the reductive TCA cycle rather than directly to pyruvate. This redistribution appears to be due to disturbance of the physiologically optimal ATP:NADPH ratio caused by limonene synthesis. Additionally, we find evidence of metabolite channeling in *Synechococcus* 7002 in the amphibolic loop involving PEP carboxylase, malate dehydrogenase, and the malic enzyme. This study provides a new avenue for increasing titers of terpenoid products in the amphibolic reactions by proposing ATP:NADPH ratios as an important consideration for limonene production, and providing a potential mechanism for recovery or the optimal ratio in response to the production of limonene. We also highlight the advantages of INST-^13^C-MFA as an essential analytical tool for characterizing and understanding cyanobacterial phenotypes from a whole-cell perspective.

## Acknowledgements

This work was supported by NSF grant # 1604691. We would like to thank Amy Zheng, Dr. Doug Allen and Dr. Yinjie Tang for their helpful advice and discussions about implementing INCA.

## Nomenclature

**Table 1.** Abbreviations

| Abbreviation | Long Name |
| --- | --- |
| **Metabolites** |  |
| 2PG | 2-phosphoglycolate |
| 2PGA | 2-phosphoglycerate |
| 3PG | 3-phosphoglycerate |
| ACA | acetyl-CoA |
| ADP | adenosine diphosphate |
| AKG | alpha-ketoglutarate |
| ATP | adenosine triphosphate |
| CIT | citrate |
| DHAP | dihydroxyacetone phosphate |
| DMAPP | dimethylallyl pyrophosphate |
| DOXP | deoxyxylulose 5-phosphate |
| E4P | erythrose 4-phosphate |
| F6P | fructose 6-phosphate |
| FBP | fructose 1,6-bisphosphate |
| FUM | fumarate |
| G1P | glucose 1-phosphate |
| G6P | glucose 6-phosphate |
| GAP | glyceraldehyde 3-phosphate |
| GLYC | glycolate |
| GOX | glyoxylate |
| ICI | isocitrate |
| IPP | isopentenyl pyrophosphate |
| LIM | limonene |
| MAL | malate |
| NAD<sup>+</sup> | nicotinamide adenine dinucleotide |
| NADH | nicotinamide adenine dinucleotide hydride |
| NADP<sup>+</sup> | nicotinamide adenine dinucleotide phosphate |
| NADPH | nicotinamide adenine dinucleotide phosphate hydride |
| OAA | oxaloacetate |
| PEP | phosphoenolpyruvate |
| PYR | pyruvate |
| R5P | ribose 5-phosphate |
| RU5P | ribulose 5-phosphate |
| RUBP | ribulose 1,5-bisphosphate |
| S7P | sedoheptulose 7-phosphate |
| SBP | sedoheptulose 1,7-bisphosphate |
| SSA | succinic semialdehyde |
| SUC | succinate |
| X5P | xylulose 5-phosphate |
| **Enzymes** |  |
| LS | limonene synthase |
| MDH | malate dehydrogenase |
| ME | malic enzyme |
| PEPc | phosphoenolpyruvate carboxylase |
| PK | pyruvate kinase |

## References

Abernathy, M.H., Czajka, J., Allen, D.K., Hill, N.C., Cameron, C., and Tang, Y.J. (2019). Cyanobacterial carboxysome mutant analysis reveals the influence of enzyme compartmentalization on cellular metabolism and metabolic network rigidity. 54, 222–231.

Abernathy, M.H., Yu, J., Ma, F., Liberton, M., Ungerer, J., Hollinshead, W.D., Gopalakrishnan, S., He, L., Maranas, C.D., Pakrasi, H.B., Allen, D.K., and Tang, Y.J. (2017). Deciphering cyanobacterial phenotypes for fast photoautotrophic growth via isotopically nonstationary metabolic flux analysis. Biotechnology for Biofuels 10, 1–13.

Abramson, B.W., Kachel, B., Kramer, D.M., and Ducat, D.C. (2016). Increased Photochemical Efficiency in Cyanobacteria via an Engineered Sucrose Sink. Plant and Cell Physiology 57, 2451–2460.

Abramson, B.W., Lensmire, J., Lin, Y.-T., Jennings, E., and Ducat, D.C. (2018). Redirecting carbon to bioproduction via a growth arrest switch in a sucrose-secreting cyanobacterium. Algal Research 33, 248–255.

Adebiyi, A.O., Jazmin, L.J., and Young, J.D. (2015). 13C flux analysis of cyanobacterial metabolism. Photosynthesis Research 126, 19–32.

Alric, J., Lavergne, J., and Rappaport, F. (2010). Redox and ATP control of photosynthetic cyclic electron flow in Chlamydomonas reinhardtii (I) aerobic conditions. Biochimica et Biophysica Acta (BBA) - Bioenergetics 1797, 44–51.

Batterton, J.C., and Baalen, C. (1971). Growth responses of blue-green algae to sodium chloride concentration. Archiv fr Mikrobiologie 76, 151–165.

Berla, B.M., Saha, R., Immethun, C.M., Maranas, C.D., Moon, T.S., and Pakrasi, H.B. (2013). Synthetic biology of cyanobacteria: unique challenges and opportunities. Frontiers in Microbiology 4.

Broddrick, J.T., Rubin, B.E., Welkie, D.G., Du, N., Mih, N., Diamond, S., Lee, J.J., Golden, S.S., and Palsson, B.O. (2016). Unique attributes of cyanobacterial metabolism revealed by improved genome-scale metabolic modeling and essential gene analysis. Proceedings of the National Academy of Sciences 113, E8344–E8353.

Cheah, Y.E., Xu, Y., Sacco, S.A., Babele, P.K., Zheng, A.O., Johnson, C.H., and Young, J.D. (2020). Systematic identification and elimination of flux bottlenecks in the aldehyde production pathway of Synechococcus elongatus PCC 7942. Metabolic Engineering 60, 56–65.

Colman, B. (1989). Photosynthetic carbon assimilation and the suppression of photorespiration in the cyanobacteria. Aquatic Botany 34, 211–231.

Davies, F.K., Work, V.H., Beliaev, A.S., and Posewitz, M.C. (2014). Engineering limonene and bisabolene production in wild type and a glycogen-deficient mutant of Synechococc us. 2, 1–11.

Ducat, D.C., Way, J.C., and Silver, P.A. (2011). Engineering cyanobacteria to generate high-value products. Trends in Biotechnology 29, 95–103.

Englund, E., Shabestary, K., Hudson, E.P., and Lindberg, P. (2018). Systematic overexpression study to find target enzymes enhancing production of terpenes in Synechocystis PCC 6803, using isoprene as a model compound. Metabolic Engineering 49, 164–177.

Erdrich, P., Knoop, H., Steuer, R., and Klamt, S. (2014). Cyanobacterial biofuels: new insights and strain design strategies revealed by computational modeling. Microbial cell factories 13, 1–15.

Gao, X., Gao, F., Liu, D., Zhang, H., Nie, X., and Yang, C. (2016). Engineering the methylerythritol phosphate pathway in cyanobacteria for photosynthetic isoprene production from CO 2. Energy and Environmental Science 9, 1400–1411.

Hendry, J.I., Prasannan, C., Ma, F., Möllers, K.B., Jaiswal, D., Digmurti, M., Allen, D.K., Frigaard, N.U., Dasgupta, S., and Wangikar, P.P. (2017). Rerouting of carbon flux in a glycogen mutant of cyanobacteria assessed via isotopically non-stationary 13C metabolic flux analysis. Biotechnology and Bioengineering 114, 2298–2308.

Hendry, J.I., Prasannan, C.B., Joshi, A., Dasgupta, S., and Wangikar, P.P. (2016). Metabolic model of Synechococcus sp. PCC 7002: Prediction of flux distribution and network modification for enhanced biofuel production. Bioresour Technol 213, 190–197.

Iijima, H., Watanabe, A., Sukigara, H., Iwazumi, K., Shirai, T., Kondo, A., and Osanai, T. (2021). Four-carbon dicarboxylic acid production through the reductive branch of the open cyanobacterial tricarboxylic acid cycle in Synechocystis sp. PCC 6803. Metabolic Engineering 65, 88–98.

Ishikawa, M., Tsuchiya, D., Oyama, T., Tsunaka, Y., and Morikawa, K. (2004). Structural basis for channelling mechanism of a fatty acid β-oxidation multienzyme complex. The EMBO Journal 23, 2745–2754.

Ishikawa, Y., and Kawai-Yamada, M. (2019). Physiological significance of NAD kinases in cyanobacteria. Frontiers in Plant Science 10, 847.

Jazmin, L.J., Xu, Y., Cheah, Y.E., Adebiyi, A.O., Johnson, C.H., and Young, J.D. (2017). Isotopically nonstationary 13C flux analysis of cyanobacterial isobutyraldehyde production. Metabolic engineering 42, 9–18.

Kanehisa, M. (2002). The KEGG database.

Kanehisa, M., Goto, S., Sato, Y., Kawashima, M., Furumichi, M., and Tanabe, M. (2014). Data, information, knowledge and principle: back to metabolism in KEGG. Nucleic acids research 42, D199–D205.

Khan, A.Z., Bilal, M., Mehmood, S., Sharma, A., and Iqbal, H.M.N. (2019). State-of-the-art genetic modalities to engineer cyanobacteria for sustainable biosynthesis of biofuel and fine-chemicals to meet bio-economy challenges. Life 9, 1–22.

Knoop, H., Zilliges, Y., Lockau, W., and Steuer, R. (2010). The metabolic network of Synechocystis sp. PCC 6803: systemic properties of autotrophic growth. Plant Physiol 154, 410–422.

Kramer, D.M., and Evans, J.R. (2011). The importance of energy balance in improving photosynthetic productivity. Plant physiology 155, 70–78.

Lin, P.-C., Zhang, F., and Pakrasi, H.B. (2020). Enhanced production of sucrose in the fast-growing cyanobacterium Synechococcus elongatus UTEX 2973. Scientific reports 10, 1–8.

Ludwig, M., and Bryant, D.A. (2012). Acclimation of the Global Transcriptome of the Cyanobacterium Synechococcus sp. Strain PCC 7002 to Nutrient Limitations and Different Nitrogen Sources. Frontiers in Microbiology 3.

Nakajima, T., Yoshikawa, K., Toya, Y., Matsuda, F., and Shimizu, H. (2017). Metabolic flux analysis of Synechocystis sp. PCC 6803 Δ nrtABCD mutant reveals a mechanism for metabolic adaptation to nitrogen-limited conditions. Plant and Cell Physiology, pcw233.

Negi, S., Barry, A.N., Friedland, N., Sudasinghe, N., Subramanian, S., Pieris, S., Holguin, F.O., Dungan, B., Schaub, T., and Sayre, R. (2015). Impact of nitrogen limitation on biomass, photosynthesis, and lipid accumulation in Chlorella sorokiniana. Journal of Applied Phycology 28, 803–812.

Nomura, C.T., Sakamoto, T., and Bryant, D.A. (2006). Roles for heme-copper oxidases in extreme highlight and oxidative stress response in the cyanobacterium Synechococcus sp. PCC 7002. Arch Microbiol 185, 471–479.

Qian, X., Zhang, Y., Lun, D.S., and Dismukes, G.C. (2018). Rerouting of Metabolism into Desired Cellular Products by Nutrient Stress: Fluxes Reveal the Selected Pathways in Cyanobacterial Photosynthesis. ACS Synthetic Biology 7, 1465–1476.

Sake, C.L., Newman, D.M., and Boyle, N.R. (2020). Evaluation of quenching methods for metabolite recovery in photoautotrophic Synechococcus sp. PCC 7002. Biotechnology Progress 36.

Singh, R., Lemire, J., Mailloux, R.J., and Appanna, V.D. (2008). A Novel Strategy Involved Anti-Oxidative Defense: The Conversion of NADH into NADPH by a Metabolic Network. PLoS ONE 3, e2682.

Spaans, S., Weusthuis, R., Van Der Oost, J., and Kengen, S. (2015). NADPH-generating systems in bacteria and archaea. Frontiers in Microbiology 6.

Thiel, K., Patrikainen, P., Nagy, C., Fitzpatrick, D., Pope, N., Aro, E.-M., and Kallio, P. (2019). Redirecting photosynthetic electron flux in the cyanobacterium Synechocystis sp. PCC 6803 by the deletion of flavodiiron protein Flv3. Microbial Cell Factories 18.

Trost, P., and Lemaire, S.D. (2013). Redox regulation of the Calvin – Benson cycle: something old, something new.

Wang, X., Liu, W., Xin, C., Zheng, Y., Cheng, Y., Sun, S., Li, R., Zhu, X.-G., Dai, S.Y., Rentzepis, P.M., and Yuan, J.S. (2016). Enhanced limonene production in cyanobacteria reveals photosynthesis limitations. Proceedings of the National Academy of Sciences 113, 14225–14230.

Wilde, A., and Hihara, Y. (2016). Transcriptional and posttranscriptional regulation of cyanobacterial photosynthesis. Biochimica et Biophysica Acta - Bioenergetics 1857, 296–308.

Xiong, W., Cano, M., Wang, B., and Douchi, D. (2017). The plasticity of cyanobacterial carbon metabolism. Current Opinion in Chemical Biology 41, 12–19.

You, L., Berla, B., He, L., Pakrasi, H.B., and Tang, Y.J. (2014). 13C-MFA delineates the photomixotrophic metabolism ofSynechocystissp. PCC 6803 under light- and carbon-sufficient conditions. Biotechnology Journal 9, 684–692.

Young, J.D. (2014). INCA: A computational platform for isotopically non-stationary metabolic flux analysis. Bioinformatics 30, 1333–1335.

Young, J.D., Shastri, A.A., Stephanopoulos, G., and Morgan, J.A. (2011). Mapping photoautotrophic metabolism with isotopically nonstationary (13)C flux analysis. Metab Eng 13, 656–665.

Yu King Hing, N., Liang, F., Lindblad, P., and Morgan, J.A. (2019). Combining isotopically non-stationary metabolic flux analysis with proteomics to unravel the regulation of the Calvin-Benson-Bassham cycle in Synechocystis sp. PCC 6803. Metabolic Engineering 56, 77–84.

Zhang, S., and Bryant, D.A. (2011). The tricarboxylic acid cycle in cyanobacteria. Science 334, 1551–1553.

