## Supplemental Info for "Characterizing Photosynthetic Biofuel Production: Isotopically non-stationary ^13^C metabolic flux analysis (INST-^13^CMFA) on limonene producing *Synechococcus* sp. PCC 7002"

Supplementary Information

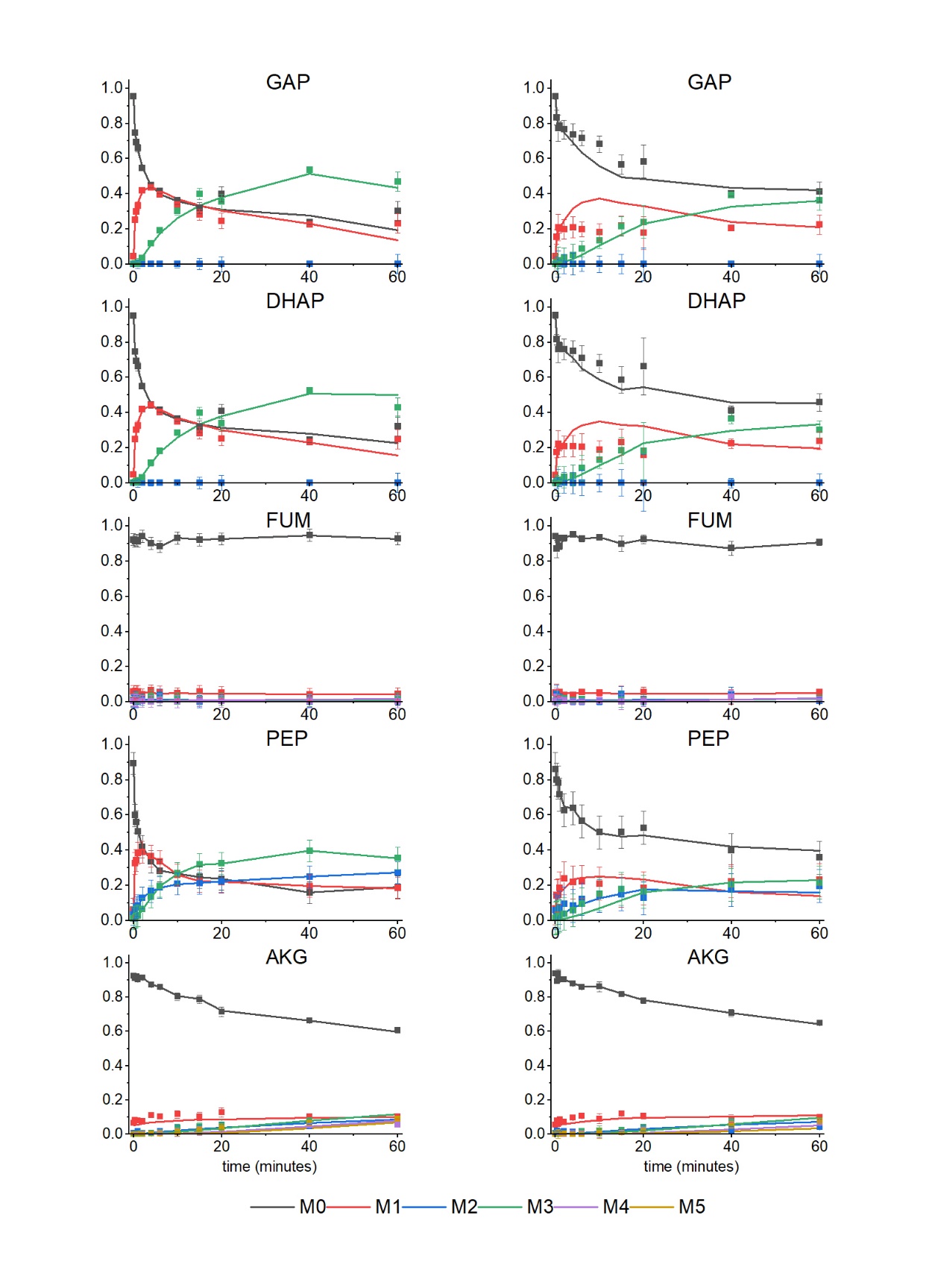
**Figure S1: Labeling dynamics of measured metabolites.** Mass isotopomer distributions of key intracellular metabolites over time are calculated as the mean of three biological replicates (n=3) and error bars represent the highest value of either standard error between replicates or to the error required to correct the zero timepoint value to the distribution vector expected due to natural ^13^C abundances.

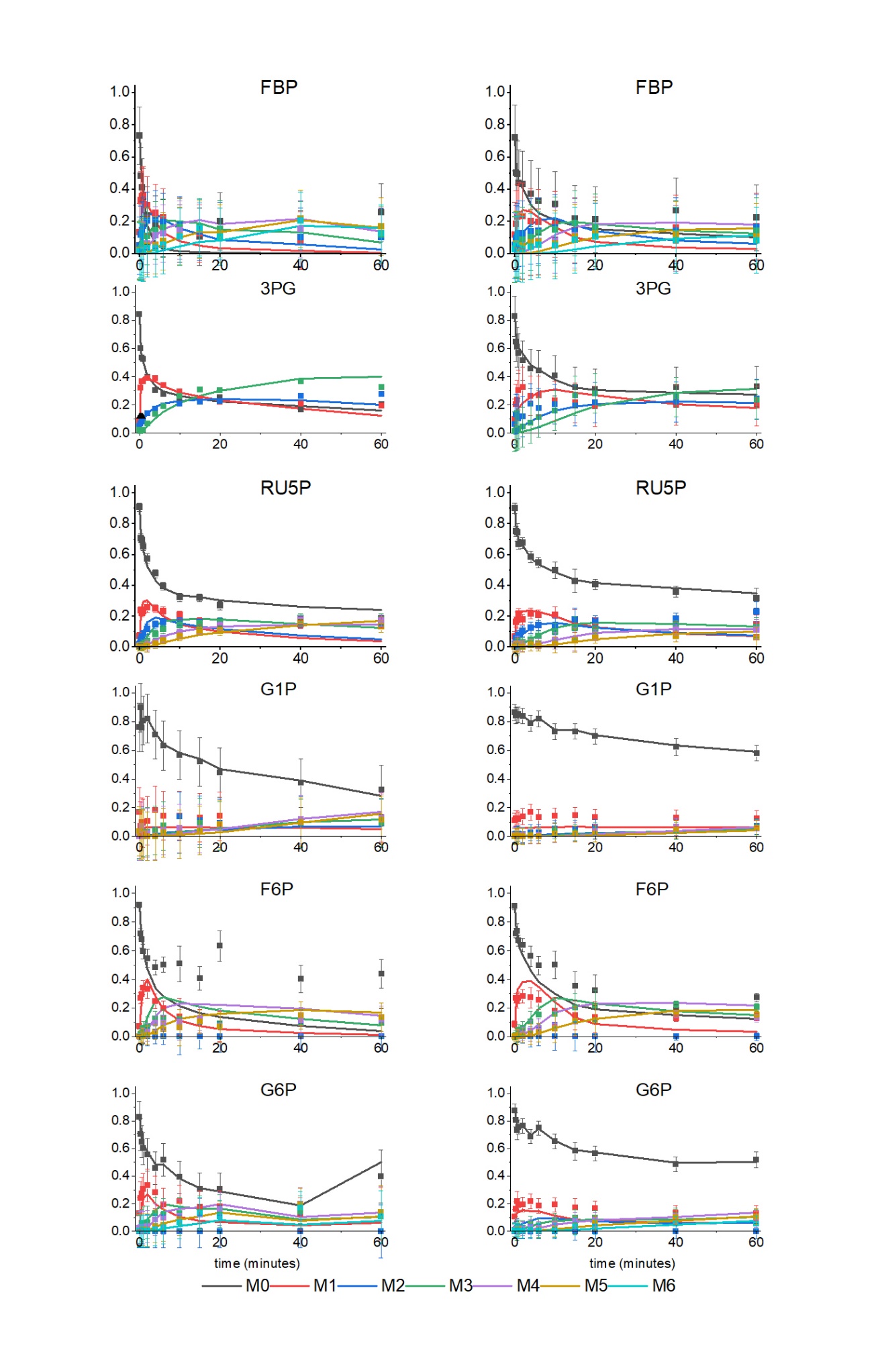

**Table S1: Relative isotopomer abundances for key measured metabolites in wild type *Synechococcus* 7002.**

| Metabolite | Isotopomer | Time (mins) | | | | | | | | | | | |
| --- | --- | --- | --- | --- | --- | --- | --- | --- | --- | --- | --- | --- | --- |
|  |  | 0 | 0.33 | 0.66 | 1 | 2 | 4 | 6 | 10 | 15 | 20 | 40 | 60 |
| 3PG | M0 | 0.8403 | 0.6007 | 0.5360 | 0.5242 | 0.3989 | 0.3028 | 0.2764 | 0.2407 | 0.2239 | 0.2508 | 0.1648 | 0.2051 |
|  | M1 | 0.0892 | 0.3224 | 0.3658 | 0.3657 | 0.3906 | 0.3908 | 0.3410 | 0.2915 | 0.2437 | 0.2206 | 0.2079 | 0.1951 |
|  | M2 | 0.0502 | 0.0660 | 0.0800 | 0.0886 | 0.1416 | 0.1728 | 0.1926 | 0.2068 | 0.2229 | 0.2252 | 0.2613 | 0.2740 |
|  | M3 | 0.0203 | 0.0109 | 0.0182 | 0.0215 | 0.0688 | 0.1336 | 0.1900 | 0.2610 | 0.3095 | 0.3033 | 0.3660 | 0.3258 |
|  | M0 error | 0.1176 | 0.1176 | 0.1176 | 0.1176 | 0.1176 | 0.1176 | 0.1176 | 0.1176 | 0.1176 | 0.1176 | 0.1176 | 0.1176 |
|  | M1 error | 0.1176 | 0.1176 | 0.1176 | 0.1176 | 0.1176 | 0.1176 | 0.1176 | 0.1176 | 0.1176 | 0.1176 | 0.1176 | 0.1176 |
|  | M2 error | 0.1176 | 0.1176 | 0.1176 | 0.1176 | 0.1176 | 0.1176 | 0.1176 | 0.1176 | 0.1176 | 0.1176 | 0.1176 | 0.1176 |
|  | M3 error | 0.1176 | 0.1176 | 0.1176 | 0.1176 | 0.1176 | 0.1176 | 0.1176 | 0.1176 | 0.1176 | 0.1176 | 0.1176 | 0.1176 |
| G6P | M0 | 0.8269 | 0.7013 | 0.6490 | 0.6002 | 0.5562 | 0.4587 | 0.5177 | 0.3912 | 0.3067 | 0.3047 | 0.1715 | 0.3984 |
|  | M1 | 0.1270 | 0.2400 | 0.2744 | 0.3032 | 0.3327 | 0.2824 | 0.1955 | 0.2184 | 0.1751 | 0.1779 | 0.1237 | 0.1253 |
|  | M2 | 0.0000 | 0.0000 | 0.0000 | 0.0000 | 0.0000 | 0.0000 | 0.0000 | 0.0000 | 0.0000 | 0.0000 | 0.0000 | 0.0000 |
|  | M3 | 0.0166 | 0.0280 | 0.0421 | 0.0581 | 0.0646 | 0.1281 | 0.1213 | 0.1294 | 0.1402 | 0.1464 | 0.1409 | 0.1033 |
|  | M4 | 0.0225 | 0.0272 | 0.0312 | 0.0263 | 0.0344 | 0.0837 | 0.0944 | 0.1118 | 0.1195 | 0.1616 | 0.1983 | 0.1326 |
|  | M5 | 0.0048 | 0.0020 | 0.0016 | 0.0106 | 0.0087 | 0.0357 | 0.0439 | 0.0847 | 0.1310 | 0.1032 | 0.1936 | 0.1366 |
|  | M6 | 0.0023 | 0.0015 | 0.0016 | 0.0016 | 0.0034 | 0.0113 | 0.0272 | 0.0645 | 0.1276 | 0.1062 | 0.1720 | 0.1038 |
|  | M0 error | 0.1171 | 0.1171 | 0.1171 | 0.1171 | 0.1171 | 0.1171 | 0.1171 | 0.1171 | 0.1171 | 0.1171 | 0.1171 | 0.1916 |
|  | M1 error | 0.1171 | 0.1171 | 0.1171 | 0.1171 | 0.1171 | 0.1171 | 0.1171 | 0.1171 | 0.1171 | 0.1171 | 0.1171 | 0.1916 |
|  | M2 error | 0.1171 | 0.1171 | 0.1171 | 0.1171 | 0.1171 | 0.1171 | 0.1171 | 0.1171 | 0.1171 | 0.1171 | 0.1171 | 0.1916 |
|  | M3 error | 0.1171 | 0.1171 | 0.1171 | 0.1171 | 0.1171 | 0.1171 | 0.1171 | 0.1171 | 0.1171 | 0.1171 | 0.1171 | 0.1916 |
|  | M4 error | 0.1171 | 0.1171 | 0.1171 | 0.1171 | 0.1171 | 0.1171 | 0.1171 | 0.1171 | 0.1171 | 0.1171 | 0.1171 | 0.1916 |
|  | M5 error | 0.1171 | 0.1171 | 0.1171 | 0.1171 | 0.1171 | 0.1171 | 0.1171 | 0.1171 | 0.1171 | 0.1171 | 0.1171 | 0.1916 |
|  | M6 error | 0.1171 | 0.1171 | 0.1171 | 0.1171 | 0.1171 | 0.1171 | 0.1171 | 0.1171 | 0.1171 | 0.1171 | 0.1171 | 0.1916 |
| AKG | M0 | 0.9269 | 0.9117 | 0.9207 | 0.9032 | 0.9133 | 0.8735 | 0.8586 | 0.8039 | 0.7878 | 0.7141 | 0.6625 | 0.6071 |
|  | M1 | 0.0635 | 0.0801 | 0.0767 | 0.0798 | 0.0758 | 0.1105 | 0.1032 | 0.1151 | 0.1034 | 0.1254 | 0.1023 | 0.1033 |
|  | M2 | 0.0051 | 0.0054 | 0.0000 | 0.0148 | 0.0050 | 0.0068 | 0.0172 | 0.0254 | 0.0295 | 0.0378 | 0.0454 | 0.0548 |
|  | M3 | 0.0024 | 0.0021 | 0.0021 | 0.0013 | 0.0050 | 0.0058 | 0.0139 | 0.0368 | 0.0442 | 0.0548 | 0.0772 | 0.0900 |
|  | M4 | 0.0014 | 0.0007 | 0.0000 | 0.0006 | 0.0009 | 0.0015 | 0.0058 | 0.0088 | 0.0157 | 0.0328 | 0.0480 | 0.0557 |
|  | M5 | 0.0007 | 0.0000 | 0.0005 | 0.0003 | 0.0000 | 0.0018 | 0.0012 | 0.0100 | 0.0194 | 0.0350 | 0.0646 | 0.0891 |
|  | M0 error | 0.0112 | 0.0128 | 0.0112 | 0.0112 | 0.0112 | 0.0128 | 0.0112 | 0.0222 | 0.0235 | 0.0270 | 0.0124 | 0.0154 |
|  | M1 error | 0.0112 | 0.0128 | 0.0112 | 0.0112 | 0.0112 | 0.0128 | 0.0112 | 0.0222 | 0.0235 | 0.0270 | 0.0124 | 0.0154 |
|  | M2 error | 0.0112 | 0.0128 | 0.0112 | 0.0112 | 0.0112 | 0.0128 | 0.0112 | 0.0222 | 0.0235 | 0.0270 | 0.0124 | 0.0154 |
|  | M3 error | 0.0112 | 0.0128 | 0.0112 | 0.0112 | 0.0112 | 0.0128 | 0.0112 | 0.0222 | 0.0235 | 0.0270 | 0.0124 | 0.0154 |
|  | M4 error | 0.0112 | 0.0128 | 0.0112 | 0.0112 | 0.0112 | 0.0128 | 0.0112 | 0.0222 | 0.0235 | 0.0270 | 0.0124 | 0.0154 |
|  | M5 error | 0.0112 | 0.0128 | 0.0112 | 0.0112 | 0.0112 | 0.0128 | 0.0112 | 0.0222 | 0.0235 | 0.0270 | 0.0124 | 0.0154 |
| DHAP | M0 | 0.9520 | 0.7478 | 0.6933 | 0.6633 | 0.5488 | 0.4457 | 0.4165 | 0.3640 | 0.3193 | 0.4072 | 0.2449 | 0.3233 |
|  | M1 | 0.0462 | 0.2489 | 0.3008 | 0.3268 | 0.4188 | 0.4410 | 0.4021 | 0.3492 | 0.2828 | 0.2529 | 0.2308 | 0.2474 |
|  | M2 | 0.0000 | 0.0000 | 0.0000 | 0.0000 | 0.0000 | 0.0000 | 0.0000 | 0.0000 | 0.0000 | 0.0000 | 0.0000 | 0.0000 |
|  | M3 | 0.0017 | 0.0033 | 0.0059 | 0.0099 | 0.0324 | 0.1134 | 0.1814 | 0.2867 | 0.3979 | 0.3399 | 0.5243 | 0.4293 |
|  | M0 error | 0.0135 | 0.0135 | 0.0135 | 0.0241 | 0.0135 | 0.0174 | 0.0135 | 0.0154 | 0.0328 | 0.0411 | 0.0156 | 0.0554 |
|  | M1 error | 0.0135 | 0.0135 | 0.0135 | 0.0241 | 0.0135 | 0.0174 | 0.0135 | 0.0154 | 0.0328 | 0.0411 | 0.0156 | 0.0554 |
|  | M2 error | 0.0135 | 0.0135 | 0.0135 | 0.0241 | 0.0135 | 0.0174 | 0.0135 | 0.0154 | 0.0328 | 0.0411 | 0.0156 | 0.0554 |
|  | M3 error | 0.0135 | 0.0135 | 0.0135 | 0.0241 | 0.0135 | 0.0174 | 0.0135 | 0.0154 | 0.0328 | 0.0411 | 0.0156 | 0.0554 |
| F6P | M0 | 0.9181 | 0.7169 | 0.6788 | 0.5942 | 0.5460 | 0.4829 | 0.5018 | 0.5072 | 0.4063 | 0.6324 | 0.4020 | 0.4387 |
|  | M1 | 0.0721 | 0.2673 | 0.2932 | 0.3416 | 0.3314 | 0.2474 | 0.1960 | 0.1389 | 0.1088 | 0.0710 | 0.0884 | 0.0953 |
|  | M2 | 0.0000 | 0.0000 | 0.0000 | 0.0000 | 0.0000 | 0.0000 | 0.0000 | 0.0000 | 0.0000 | 0.0000 | 0.0000 | 0.0000 |
|  | M3 | 0.0027 | 0.0131 | 0.0227 | 0.0543 | 0.0737 | 0.1277 | 0.1318 | 0.1218 | 0.1250 | 0.0897 | 0.0965 | 0.0967 |
|  | M4 | 0.0028 | 0.0027 | 0.0028 | 0.0064 | 0.0310 | 0.0938 | 0.0965 | 0.1041 | 0.1263 | 0.0727 | 0.1075 | 0.1130 |
|  | M5 | 0.0011 | 0.0000 | 0.0025 | 0.0024 | 0.0103 | 0.0318 | 0.0480 | 0.0639 | 0.1046 | 0.0667 | 0.1489 | 0.1359 |
|  | M6 | 0.0032 | 0.0000 | 0.0000 | 0.0011 | 0.0076 | 0.0163 | 0.0258 | 0.0641 | 0.1290 | 0.0673 | 0.1566 | 0.1205 |
|  | M0 error | 0.0185 | 0.0377 | 0.0583 | 0.0497 | 0.0635 | 0.0512 | 0.0544 | 0.1266 | 0.0837 | 0.1068 | 0.0947 | 0.1010 |
|  | M1 error | 0.0185 | 0.0377 | 0.0583 | 0.0497 | 0.0635 | 0.0512 | 0.0544 | 0.1266 | 0.0837 | 0.1068 | 0.0947 | 0.1010 |
|  | M2 error | 0.0185 | 0.0377 | 0.0583 | 0.0497 | 0.0635 | 0.0512 | 0.0544 | 0.1266 | 0.0837 | 0.1068 | 0.0947 | 0.1010 |
|  | M3 error | 0.0185 | 0.0377 | 0.0583 | 0.0497 | 0.0635 | 0.0512 | 0.0544 | 0.1266 | 0.0837 | 0.1068 | 0.0947 | 0.1010 |
|  | M4 error | 0.0185 | 0.0377 | 0.0583 | 0.0497 | 0.0635 | 0.0512 | 0.0544 | 0.1266 | 0.0837 | 0.1068 | 0.0947 | 0.1010 |
|  | M5 error | 0.0185 | 0.0377 | 0.0583 | 0.0497 | 0.0635 | 0.0512 | 0.0544 | 0.1266 | 0.0837 | 0.1068 | 0.0947 | 0.1010 |
|  | M6 error | 0.0185 | 0.0377 | 0.0583 | 0.0497 | 0.0635 | 0.0512 | 0.0544 | 0.1266 | 0.0837 | 0.1068 | 0.0947 | 0.1010 |
| FBP | M0 | 0.7349 | 0.4850 | 0.4117 | 0.3535 | 0.2365 | 0.1822 | 0.1565 | 0.1262 | 0.1045 | 0.2029 | 0.0919 | 0.2552 |
|  | M1 | 0.1329 | 0.3307 | 0.3565 | 0.3644 | 0.2995 | 0.2530 | 0.2265 | 0.1691 | 0.1277 | 0.1405 | 0.0808 | 0.1043 |
|  | M2 | 0.0508 | 0.1169 | 0.1452 | 0.1721 | 0.2058 | 0.2121 | 0.1975 | 0.1771 | 0.1367 | 0.1421 | 0.1041 | 0.1034 |
|  | M3 | 0.0244 | 0.0296 | 0.0414 | 0.0641 | 0.1105 | 0.1459 | 0.1629 | 0.1662 | 0.1664 | 0.1548 | 0.1454 | 0.1176 |
|  | M4 | 0.0205 | 0.0151 | 0.0188 | 0.0225 | 0.0635 | 0.1159 | 0.1243 | 0.1478 | 0.1565 | 0.1254 | 0.1556 | 0.1241 |
|  | M5 | 0.0206 | 0.0126 | 0.0161 | 0.0144 | 0.0502 | 0.0554 | 0.0783 | 0.1093 | 0.1514 | 0.1268 | 0.2165 | 0.1690 |
|  | M6 | 0.0159 | 0.0100 | 0.0101 | 0.0090 | 0.0340 | 0.0354 | 0.0539 | 0.1044 | 0.1568 | 0.1075 | 0.2056 | 0.1264 |
|  | M0 error | 0.1788 | 0.1788 | 0.1788 | 0.1788 | 0.1788 | 0.1788 | 0.1788 | 0.1788 | 0.1788 | 0.1788 | 0.1788 | 0.1788 |
|  | M1 error | 0.1788 | 0.1788 | 0.1788 | 0.1788 | 0.1788 | 0.1788 | 0.1788 | 0.1788 | 0.1788 | 0.1788 | 0.1788 | 0.1788 |
|  | M2 error | 0.1788 | 0.1788 | 0.1788 | 0.1788 | 0.1788 | 0.1788 | 0.1788 | 0.1788 | 0.1788 | 0.1788 | 0.1788 | 0.1788 |
|  | M3 error | 0.1788 | 0.1788 | 0.1788 | 0.1788 | 0.1788 | 0.1788 | 0.1788 | 0.1788 | 0.1788 | 0.1788 | 0.1788 | 0.1788 |
|  | M4 error | 0.1788 | 0.1788 | 0.1788 | 0.1788 | 0.1788 | 0.1788 | 0.1788 | 0.1788 | 0.1788 | 0.1788 | 0.1788 | 0.1788 |
|  | M5 error | 0.1788 | 0.1788 | 0.1788 | 0.1788 | 0.1788 | 0.1788 | 0.1788 | 0.1788 | 0.1788 | 0.1788 | 0.1788 | 0.1788 |
|  | M6 error | 0.1788 | 0.1788 | 0.1788 | 0.1788 | 0.1788 | 0.1788 | 0.1788 | 0.1788 | 0.1788 | 0.1788 | 0.1788 | 0.1788 |
| FUM | M0 | 0.9233 | 0.9132 | 0.9203 | 0.9132 | 0.9429 | 0.9020 | 0.8837 | 0.9329 | 0.9219 | 0.9276 | 0.9480 | 0.9281 |
|  | M1 | 0.0580 | 0.0450 | 0.0542 | 0.0583 | 0.0378 | 0.0643 | 0.0547 | 0.0465 | 0.0569 | 0.0518 | 0.0409 | 0.0435 |
|  | M2 | 0.0032 | 0.0247 | 0.0000 | 0.0024 | 0.0123 | 0.0018 | 0.0390 | 0.0000 | 0.0000 | 0.0043 | 0.0000 | 0.0126 |
|  | M3 | 0.0103 | 0.0060 | 0.0176 | 0.0223 | 0.0042 | 0.0301 | 0.0134 | 0.0195 | 0.0139 | 0.0137 | 0.0071 | 0.0157 |
|  | M4 | 0.0052 | 0.0111 | 0.0080 | 0.0037 | 0.0027 | 0.0018 | 0.0092 | 0.0011 | 0.0073 | 0.0027 | 0.0040 | 0.0000 |
|  | M0 error | 0.0344 | 0.0344 | 0.0344 | 0.0344 | 0.0344 | 0.0344 | 0.0344 | 0.0344 | 0.0344 | 0.0344 | 0.0344 | 0.0344 |
|  | M1 error | 0.0344 | 0.0344 | 0.0344 | 0.0344 | 0.0344 | 0.0344 | 0.0344 | 0.0344 | 0.0344 | 0.0344 | 0.0344 | 0.0344 |
|  | M2 error | 0.0344 | 0.0344 | 0.0344 | 0.0344 | 0.0344 | 0.0344 | 0.0344 | 0.0344 | 0.0344 | 0.0344 | 0.0344 | 0.0344 |
|  | M3 error | 0.0344 | 0.0344 | 0.0344 | 0.0344 | 0.0344 | 0.0344 | 0.0344 | 0.0344 | 0.0344 | 0.0344 | 0.0344 | 0.0344 |
|  | M4 error | 0.0344 | 0.0344 | 0.0344 | 0.0344 | 0.0344 | 0.0344 | 0.0344 | 0.0344 | 0.0344 | 0.0344 | 0.0344 | 0.0344 |
| G1P | M0 | 0.7596 | 0.8980 | 0.7577 | 0.8080 | 0.8202 | 0.7097 | 0.6317 | 0.5655 | 0.5193 | 0.4450 | 0.3720 | 0.3247 |
|  | M1 | 0.1689 | 0.0721 | 0.0983 | 0.0906 | 0.1075 | 0.1805 | 0.1426 | 0.1432 | 0.1303 | 0.1410 | 0.1169 | 0.1262 |
|  | M2 | 0.0119 | 0.0000 | 0.0344 | 0.0183 | 0.0019 | 0.0420 | 0.0731 | 0.1384 | 0.1121 | 0.0910 | 0.1074 | 0.0898 |
|  | M3 | 0.0158 | 0.0000 | 0.0000 | 0.0293 | 0.0308 | 0.0402 | 0.0734 | 0.0527 | 0.0928 | 0.0711 | 0.0907 | 0.0922 |
|  | M4 | 0.0007 | 0.0000 | 0.0689 | 0.0095 | 0.0216 | 0.0042 | 0.0426 | 0.0557 | 0.0562 | 0.0812 | 0.1189 | 0.1342 |
|  | M5 | 0.0345 | 0.0106 | 0.0107 | 0.0193 | 0.0000 | 0.0000 | 0.0253 | 0.0156 | 0.0406 | 0.0817 | 0.0977 | 0.1132 |
|  | M6 | 0.0086 | 0.0193 | 0.0299 | 0.0250 | 0.0179 | 0.0234 | 0.0114 | 0.0288 | 0.0486 | 0.0890 | 0.0963 | 0.1197 |
|  | M0 error | 0.1698 | 0.1698 | 0.1698 | 0.1698 | 0.1698 | 0.1698 | 0.1698 | 0.1698 | 0.1698 | 0.1698 | 0.1698 | 0.1698 |
|  | M1 error | 0.1698 | 0.1698 | 0.1698 | 0.1698 | 0.1698 | 0.1698 | 0.1698 | 0.1698 | 0.1698 | 0.1698 | 0.1698 | 0.1698 |
|  | M2 error | 0.1698 | 0.1698 | 0.1698 | 0.1698 | 0.1698 | 0.1698 | 0.1698 | 0.1698 | 0.1698 | 0.1698 | 0.1698 | 0.1698 |
|  | M3 error | 0.1698 | 0.1698 | 0.1698 | 0.1698 | 0.1698 | 0.1698 | 0.1698 | 0.1698 | 0.1698 | 0.1698 | 0.1698 | 0.1698 |
|  | M4 error | 0.1698 | 0.1698 | 0.1698 | 0.1698 | 0.1698 | 0.1698 | 0.1698 | 0.1698 | 0.1698 | 0.1698 | 0.1698 | 0.1698 |
|  | M5 error | 0.1698 | 0.1698 | 0.1698 | 0.1698 | 0.1698 | 0.1698 | 0.1698 | 0.1698 | 0.1698 | 0.1698 | 0.1698 | 0.1698 |
|  | M6 error | 0.1698 | 0.1698 | 0.1698 | 0.1698 | 0.1698 | 0.1698 | 0.1698 | 0.1698 | 0.1698 | 0.1698 | 0.1698 | 0.1698 |
| GAP | M0 | 0.9525 | 0.7466 | 0.6947 | 0.6592 | 0.5470 | 0.4483 | 0.4151 | 0.3607 | 0.3187 | 0.3975 | 0.2394 | 0.3026 |
|  | M1 | 0.0457 | 0.2500 | 0.2997 | 0.3324 | 0.4192 | 0.4341 | 0.3942 | 0.3384 | 0.2819 | 0.2466 | 0.2240 | 0.2299 |
|  | M2 | 0.0000 | 0.0000 | 0.0000 | 0.0000 | 0.0000 | 0.0000 | 0.0000 | 0.0000 | 0.0000 | 0.0000 | 0.0000 | 0.0000 |
|  | M3 | 0.0018 | 0.0035 | 0.0056 | 0.0084 | 0.0338 | 0.1175 | 0.1907 | 0.3009 | 0.3994 | 0.3559 | 0.5366 | 0.4675 |
|  | M0 error | 0.0133 | 0.0133 | 0.0133 | 0.0226 | 0.0133 | 0.0133 | 0.0133 | 0.0194 | 0.0318 | 0.0426 | 0.0142 | 0.0552 |
|  | M1 error | 0.0133 | 0.0133 | 0.0133 | 0.0226 | 0.0133 | 0.0133 | 0.0133 | 0.0194 | 0.0318 | 0.0426 | 0.0142 | 0.0552 |
|  | M2 error | 0.0133 | 0.0133 | 0.0133 | 0.0226 | 0.0133 | 0.0133 | 0.0133 | 0.0194 | 0.0318 | 0.0426 | 0.0142 | 0.0552 |
|  | M3 error | 0.0133 | 0.0133 | 0.0133 | 0.0226 | 0.0133 | 0.0133 | 0.0133 | 0.0194 | 0.0318 | 0.0426 | 0.0142 | 0.0552 |
| PEP | M0 | 0.8933 | 0.5982 | 0.5606 | 0.5064 | 0.4201 | 0.3339 | 0.2829 | 0.2669 | 0.2471 | 0.2350 | 0.1590 | 0.1895 |
|  | M1 | 0.0626 | 0.3261 | 0.3417 | 0.3818 | 0.3883 | 0.3661 | 0.3342 | 0.2585 | 0.2226 | 0.2186 | 0.1959 | 0.1844 |
|  | M2 | 0.0338 | 0.0587 | 0.0747 | 0.0843 | 0.1283 | 0.1664 | 0.1880 | 0.2073 | 0.2120 | 0.2216 | 0.2488 | 0.2722 |
|  | M3 | 0.0103 | 0.0170 | 0.0230 | 0.0276 | 0.0633 | 0.1336 | 0.1949 | 0.2673 | 0.3183 | 0.3248 | 0.3964 | 0.3539 |
|  | M0 error | 0.0625 | 0.0625 | 0.0625 | 0.0625 | 0.0625 | 0.0625 | 0.0625 | 0.0625 | 0.0625 | 0.0625 | 0.0625 | 0.0625 |
|  | M1 error | 0.0625 | 0.0625 | 0.0625 | 0.0625 | 0.0625 | 0.0625 | 0.0625 | 0.0625 | 0.0625 | 0.0625 | 0.0625 | 0.0625 |
|  | M2 error | 0.0625 | 0.0625 | 0.0625 | 0.0625 | 0.0625 | 0.0625 | 0.0625 | 0.0625 | 0.0625 | 0.0625 | 0.0625 | 0.0625 |
|  | M3 error | 0.0625 | 0.0625 | 0.0625 | 0.0625 | 0.0625 | 0.0625 | 0.0625 | 0.0625 | 0.0625 | 0.0625 | 0.0625 | 0.0625 |
| R5P | M0 | 0.9081 | 0.7084 | 0.6926 | 0.6540 | 0.5722 | 0.4773 | 0.3947 | 0.3229 | 0.3213 | 0.2690 | 0.1731 | 0.1820 |
|  | M1 | 0.0696 | 0.2359 | 0.2229 | 0.2353 | 0.2581 | 0.2492 | 0.2347 | 0.2078 | 0.1680 | 0.1663 | 0.1480 | 0.1599 |
|  | M2 | 0.0169 | 0.0482 | 0.0730 | 0.0835 | 0.1156 | 0.1448 | 0.1628 | 0.1680 | 0.1568 | 0.1603 | 0.1535 | 0.1741 |
|  | M3 | 0.0016 | 0.0044 | 0.0063 | 0.0222 | 0.0395 | 0.0832 | 0.1152 | 0.1436 | 0.1484 | 0.1642 | 0.1896 | 0.1787 |
|  | M4 | 0.0027 | 0.0011 | 0.0017 | 0.0037 | 0.0115 | 0.0286 | 0.0552 | 0.0912 | 0.1023 | 0.1366 | 0.1779 | 0.1722 |
|  | M5 | 0.0011 | 0.0022 | 0.0035 | 0.0013 | 0.0031 | 0.0168 | 0.0375 | 0.0666 | 0.1032 | 0.1035 | 0.1580 | 0.1332 |
|  | M0 error | 0.0260 | 0.0260 | 0.0260 | 0.0260 | 0.0334 | 0.0260 | 0.0260 | 0.0260 | 0.0260 | 0.0260 | 0.0260 | 0.0363 |
|  | M1 error | 0.0260 | 0.0260 | 0.0260 | 0.0260 | 0.0334 | 0.0260 | 0.0260 | 0.0260 | 0.0260 | 0.0260 | 0.0260 | 0.0363 |
|  | M2 error | 0.0260 | 0.0260 | 0.0260 | 0.0260 | 0.0334 | 0.0260 | 0.0260 | 0.0260 | 0.0260 | 0.0260 | 0.0260 | 0.0363 |
|  | M3 error | 0.0260 | 0.0260 | 0.0260 | 0.0260 | 0.0334 | 0.0260 | 0.0260 | 0.0260 | 0.0260 | 0.0260 | 0.0260 | 0.0363 |
|  | M4 error | 0.0260 | 0.0260 | 0.0260 | 0.0260 | 0.0334 | 0.0260 | 0.0260 | 0.0260 | 0.0260 | 0.0260 | 0.0260 | 0.0363 |
|  | M5 error | 0.0260 | 0.0260 | 0.0260 | 0.0260 | 0.0334 | 0.0260 | 0.0260 | 0.0260 | 0.0260 | 0.0260 | 0.0260 | 0.0363 |

**Table S2: Relative isotopomer abundances for key measured metabolites in limonene producing *Synechococcus* 7002.**

| Metabolite | Isotopomer | Time (mins) | | | | | | | | | | | |
| --- | --- | --- | --- | --- | --- | --- | --- | --- | --- | --- | --- | --- | --- |
|  |  | 0 | 0.33 | 0.66 | 1 | 2 | 4 | 6 | 10 | 15 | 20 | 40 | 60 |
| 3PG | M0 | 0.8285 | 0.6484 | 0.6092 | 0.5645 | 0.5148 | 0.4549 | 0.4442 | 0.4077 | 0.3253 | 0.3131 | 0.3252 | 0.3293 |
|  | M1 | 0.0985 | 0.2037 | 0.2335 | 0.3031 | 0.3252 | 0.2634 | 0.2656 | 0.2290 | 0.2152 | 0.1896 | 0.1987 | 0.1925 |
|  | M2 | 0.0617 | 0.1324 | 0.1384 | 0.1028 | 0.1174 | 0.2096 | 0.1780 | 0.2042 | 0.1939 | 0.2160 | 0.2208 | 0.2415 |
|  | M3 | 0.0113 | 0.0155 | 0.0189 | 0.0296 | 0.0425 | 0.0721 | 0.1123 | 0.1591 | 0.2656 | 0.2813 | 0.2553 | 0.2367 |
|  | M0 error | 0.1414 | 0.1414 | 0.1414 | 0.1414 | 0.1414 | 0.1414 | 0.1414 | 0.1414 | 0.1414 | 0.1414 | 0.1414 | 0.1414 |
|  | M1 error | 0.1414 | 0.1414 | 0.1414 | 0.1414 | 0.1414 | 0.1414 | 0.1414 | 0.1414 | 0.1414 | 0.1414 | 0.1414 | 0.1414 |
|  | M2 error | 0.1414 | 0.1414 | 0.1414 | 0.1414 | 0.1414 | 0.1414 | 0.1414 | 0.1414 | 0.1414 | 0.1414 | 0.1414 | 0.1414 |
|  | M3 error | 0.1414 | 0.1414 | 0.1414 | 0.1414 | 0.1414 | 0.1414 | 0.1414 | 0.1414 | 0.1414 | 0.1414 | 0.1414 | 0.1414 |
| G6P | M0 | 0.8733 | 0.8070 | 0.7325 | 0.7564 | 0.7630 | 0.6869 | 0.7480 | 0.6539 | 0.5825 | 0.5649 | 0.4852 | 0.5188 |
|  | M1 | 0.1052 | 0.1623 | 0.2175 | 0.1874 | 0.1937 | 0.2160 | 0.1842 | 0.1920 | 0.1727 | 0.1658 | 0.1351 | 0.1301 |
|  | M2 | 0.0000 | 0.0000 | 0.0000 | 0.0000 | 0.0000 | 0.0000 | 0.0000 | 0.0000 | 0.0000 | 0.0000 | 0.0000 | 0.0000 |
|  | M3 | 0.0104 | 0.0058 | 0.0225 | 0.0264 | 0.0209 | 0.0534 | 0.0384 | 0.0798 | 0.0942 | 0.0973 | 0.1168 | 0.1027 |
|  | M4 | 0.0047 | 0.0157 | 0.0174 | 0.0137 | 0.0192 | 0.0308 | 0.0199 | 0.0524 | 0.0718 | 0.0821 | 0.1081 | 0.0878 |
|  | M5 | 0.0041 | 0.0041 | 0.0067 | 0.0139 | 0.0018 | 0.0069 | 0.0063 | 0.0151 | 0.0419 | 0.0583 | 0.1013 | 0.1044 |
|  | M6 | 0.0024 | 0.0052 | 0.0034 | 0.0022 | 0.0015 | 0.0061 | 0.0033 | 0.0068 | 0.0369 | 0.0316 | 0.0534 | 0.0561 |
|  | M0 error | 0.0525 | 0.0525 | 0.0706 | 0.0525 | 0.0525 | 0.0525 | 0.0525 | 0.0525 | 0.0653 | 0.0525 | 0.0525 | 0.0568 |
|  | M1 error | 0.0525 | 0.0525 | 0.0706 | 0.0525 | 0.0525 | 0.0525 | 0.0525 | 0.0525 | 0.0653 | 0.0525 | 0.0525 | 0.0568 |
|  | M2 error | 0.0525 | 0.0525 | 0.0706 | 0.0525 | 0.0525 | 0.0525 | 0.0525 | 0.0525 | 0.0653 | 0.0525 | 0.0525 | 0.0568 |
|  | M3 error | 0.0525 | 0.0525 | 0.0706 | 0.0525 | 0.0525 | 0.0525 | 0.0525 | 0.0525 | 0.0653 | 0.0525 | 0.0525 | 0.0568 |
|  | M4 error | 0.0525 | 0.0525 | 0.0706 | 0.0525 | 0.0525 | 0.0525 | 0.0525 | 0.0525 | 0.0653 | 0.0525 | 0.0525 | 0.0568 |
|  | M5 error | 0.0525 | 0.0525 | 0.0706 | 0.0525 | 0.0525 | 0.0525 | 0.0525 | 0.0525 | 0.0653 | 0.0525 | 0.0525 | 0.0568 |
|  | M6 error | 0.0525 | 0.0525 | 0.0706 | 0.0525 | 0.0525 | 0.0525 | 0.0525 | 0.0525 | 0.0653 | 0.0525 | 0.0525 | 0.0568 |
| AKG | M0 | 0.9378 | 0.8951 | 0.9378 | 0.9044 | 0.9053 | 0.8804 | 0.8592 | 0.8621 | 0.8194 | 0.7808 | 0.7084 | 0.6486 |
|  | M1 | 0.0553 | 0.0787 | 0.0578 | 0.0855 | 0.0726 | 0.0947 | 0.1043 | 0.0895 | 0.1185 | 0.1055 | 0.0894 | 0.0973 |
|  | M2 | 0.0045 | 0.0207 | 0.0032 | 0.0053 | 0.0154 | 0.0121 | 0.0144 | 0.0187 | 0.0197 | 0.0347 | 0.0211 | 0.0388 |
|  | M3 | 0.0009 | 0.0031 | 0.0003 | 0.0021 | 0.0022 | 0.0072 | 0.0159 | 0.0168 | 0.0235 | 0.0413 | 0.0716 | 0.0779 |
|  | M4 | 0.0007 | 0.0015 | 0.0000 | 0.0022 | 0.0009 | 0.0054 | 0.0054 | 0.0066 | 0.0104 | 0.0218 | 0.0584 | 0.0631 |
|  | M5 | 0.0008 | 0.0009 | 0.0008 | 0.0006 | 0.0036 | 0.0002 | 0.0007 | 0.0063 | 0.0085 | 0.0158 | 0.0513 | 0.0743 |
|  | M0 error | 0.0075 | 0.0150 | 0.0266 | 0.0075 | 0.0141 | 0.0075 | 0.0083 | 0.0297 | 0.0082 | 0.0176 | 0.0222 | 0.0075 |
|  | M1 error | 0.0075 | 0.0150 | 0.0266 | 0.0075 | 0.0141 | 0.0075 | 0.0083 | 0.0297 | 0.0082 | 0.0176 | 0.0222 | 0.0075 |
|  | M2 error | 0.0075 | 0.0150 | 0.0266 | 0.0075 | 0.0141 | 0.0075 | 0.0083 | 0.0297 | 0.0082 | 0.0176 | 0.0222 | 0.0075 |
|  | M3 error | 0.0075 | 0.0150 | 0.0266 | 0.0075 | 0.0141 | 0.0075 | 0.0083 | 0.0297 | 0.0082 | 0.0176 | 0.0222 | 0.0075 |
|  | M4 error | 0.0075 | 0.0150 | 0.0266 | 0.0075 | 0.0141 | 0.0075 | 0.0083 | 0.0297 | 0.0082 | 0.0176 | 0.0222 | 0.0075 |
|  | M5 error | 0.0075 | 0.0150 | 0.0266 | 0.0075 | 0.0141 | 0.0075 | 0.0083 | 0.0297 | 0.0082 | 0.0176 | 0.0222 | 0.0075 |
| DHAP | M0 | 0.9524 | 0.8152 | 0.7603 | 0.7811 | 0.7587 | 0.7491 | 0.7089 | 0.6796 | 0.5847 | 0.6632 | 0.4109 | 0.4573 |
|  | M1 | 0.0458 | 0.1733 | 0.2225 | 0.2044 | 0.2086 | 0.2092 | 0.2061 | 0.1878 | 0.2318 | 0.1568 | 0.2239 | 0.2398 |
|  | M2 | 0.0000 | 0.0000 | 0.0000 | 0.0000 | 0.0000 | 0.0000 | 0.0000 | 0.0000 | 0.0000 | 0.0000 | 0.0000 | 0.0000 |
|  | M3 | 0.0018 | 0.0116 | 0.0172 | 0.0144 | 0.0328 | 0.0417 | 0.0850 | 0.1326 | 0.1836 | 0.1800 | 0.3653 | 0.3029 |
|  | M0 error | 0.0172 | 0.0308 | 0.0747 | 0.0210 | 0.0588 | 0.0604 | 0.0725 | 0.0508 | 0.0753 | 0.1622 | 0.0273 | 0.0510 |
|  | M1 error | 0.0172 | 0.0308 | 0.0747 | 0.0210 | 0.0588 | 0.0604 | 0.0725 | 0.0508 | 0.0753 | 0.1622 | 0.0273 | 0.0510 |
|  | M2 error | 0.0172 | 0.0308 | 0.0747 | 0.0210 | 0.0588 | 0.0604 | 0.0725 | 0.0508 | 0.0753 | 0.1622 | 0.0273 | 0.0510 |
|  | M3 error | 0.0172 | 0.0308 | 0.0747 | 0.0210 | 0.0588 | 0.0604 | 0.0725 | 0.0508 | 0.0753 | 0.1622 | 0.0273 | 0.0510 |
| F6P | M0 | 0.9100 | 0.7197 | 0.7360 | 0.6709 | 0.6376 | 0.5609 | 0.4953 | 0.4984 | 0.3547 | 0.3241 | 0.1975 | 0.2733 |
|  | M1 | 0.0871 | 0.2682 | 0.2511 | 0.2703 | 0.2820 | 0.2719 | 0.2561 | 0.1805 | 0.1487 | 0.1370 | 0.1272 | 0.1384 |
|  | M2 | 0.0000 | 0.0000 | 0.0000 | 0.0000 | 0.0000 | 0.0000 | 0.0000 | 0.0000 | 0.0000 | 0.0000 | 0.0000 | 0.0000 |
|  | M3 | 0.0022 | 0.0112 | 0.0120 | 0.0438 | 0.0582 | 0.1052 | 0.1506 | 0.1578 | 0.2037 | 0.2144 | 0.2243 | 0.2057 |
|  | M4 | 0.0000 | 0.0000 | 0.0000 | 0.0088 | 0.0174 | 0.0408 | 0.0606 | 0.0827 | 0.1014 | 0.1123 | 0.1563 | 0.1269 |
|  | M5 | 0.0008 | 0.0003 | 0.0000 | 0.0059 | 0.0048 | 0.0176 | 0.0270 | 0.0546 | 0.1091 | 0.1145 | 0.1653 | 0.1534 |
|  | M6 | 0.0000 | 0.0006 | 0.0008 | 0.0003 | 0.0000 | 0.0036 | 0.0104 | 0.0260 | 0.0825 | 0.0976 | 0.1293 | 0.1023 |
|  | M0 error | 0.0225 | 0.0319 | 0.0525 | 0.0389 | 0.0466 | 0.0721 | 0.0636 | 0.0958 | 0.0997 | 0.1088 | 0.0225 | 0.0271 |
|  | M1 error | 0.0225 | 0.0319 | 0.0525 | 0.0389 | 0.0466 | 0.0721 | 0.0636 | 0.0958 | 0.0997 | 0.1088 | 0.0225 | 0.0271 |
|  | M2 error | 0.0225 | 0.0319 | 0.0525 | 0.0389 | 0.0466 | 0.0721 | 0.0636 | 0.0958 | 0.0997 | 0.1088 | 0.0225 | 0.0271 |
|  | M3 error | 0.0225 | 0.0319 | 0.0525 | 0.0389 | 0.0466 | 0.0721 | 0.0636 | 0.0958 | 0.0997 | 0.1088 | 0.0225 | 0.0271 |
|  | M4 error | 0.0225 | 0.0319 | 0.0525 | 0.0389 | 0.0466 | 0.0721 | 0.0636 | 0.0958 | 0.0997 | 0.1088 | 0.0225 | 0.0271 |
|  | M5 error | 0.0225 | 0.0319 | 0.0525 | 0.0389 | 0.0466 | 0.0721 | 0.0636 | 0.0958 | 0.0997 | 0.1088 | 0.0225 | 0.0271 |
|  | M6 error | 0.0225 | 0.0319 | 0.0525 | 0.0389 | 0.0466 | 0.0721 | 0.0636 | 0.0958 | 0.0997 | 0.1088 | 0.0225 | 0.0271 |
| FBP | M0 | 0.7204 | 0.5035 | 0.4969 | 0.4406 | 0.4332 | 0.3737 | 0.3267 | 0.3070 | 0.2179 | 0.2123 | 0.2689 | 0.2247 |
|  | M1 | 0.1180 | 0.1872 | 0.2278 | 0.2412 | 0.2344 | 0.2016 | 0.1956 | 0.1850 | 0.1459 | 0.1469 | 0.1611 | 0.1702 |
|  | M2 | 0.0531 | 0.0959 | 0.1080 | 0.1245 | 0.1259 | 0.1392 | 0.1427 | 0.1623 | 0.1418 | 0.1247 | 0.1432 | 0.1648 |
|  | M3 | 0.0274 | 0.0553 | 0.0493 | 0.0604 | 0.0855 | 0.1003 | 0.1298 | 0.1488 | 0.1896 | 0.1676 | 0.1280 | 0.1434 |
|  | M4 | 0.0258 | 0.0465 | 0.0401 | 0.0509 | 0.0488 | 0.0694 | 0.0769 | 0.0845 | 0.1148 | 0.1135 | 0.0959 | 0.1075 |
|  | M5 | 0.0302 | 0.0606 | 0.0470 | 0.0490 | 0.0452 | 0.0719 | 0.0729 | 0.0634 | 0.1046 | 0.1276 | 0.1186 | 0.1067 |
|  | M6 | 0.0250 | 0.0509 | 0.0308 | 0.0334 | 0.0271 | 0.0439 | 0.0554 | 0.0492 | 0.0853 | 0.1075 | 0.0843 | 0.0827 |
|  | M0 error | 0.2042 | 0.2042 | 0.2042 | 0.2042 | 0.2042 | 0.2042 | 0.2042 | 0.2042 | 0.2042 | 0.2042 | 0.2042 | 0.2042 |
|  | M1 error | 0.2042 | 0.2042 | 0.2042 | 0.2042 | 0.2042 | 0.2042 | 0.2042 | 0.2042 | 0.2042 | 0.2042 | 0.2042 | 0.2042 |
|  | M2 error | 0.2042 | 0.2042 | 0.2042 | 0.2042 | 0.2042 | 0.2042 | 0.2042 | 0.2042 | 0.2042 | 0.2042 | 0.2042 | 0.2042 |
|  | M3 error | 0.2042 | 0.2042 | 0.2042 | 0.2042 | 0.2042 | 0.2042 | 0.2042 | 0.2042 | 0.2042 | 0.2042 | 0.2042 | 0.2042 |
|  | M4 error | 0.2042 | 0.2042 | 0.2042 | 0.2042 | 0.2042 | 0.2042 | 0.2042 | 0.2042 | 0.2042 | 0.2042 | 0.2042 | 0.2042 |
|  | M5 error | 0.2042 | 0.2042 | 0.2042 | 0.2042 | 0.2042 | 0.2042 | 0.2042 | 0.2042 | 0.2042 | 0.2042 | 0.2042 | 0.2042 |
|  | M6 error | 0.2042 | 0.2042 | 0.2042 | 0.2042 | 0.2042 | 0.2042 | 0.2042 | 0.2042 | 0.2042 | 0.2042 | 0.2042 | 0.2042 |
| FUM | M0 | 0.9433 | 0.8704 | 0.9321 | 0.8856 | 0.9315 | 0.9533 | 0.9276 | 0.9366 | 0.9001 | 0.9250 | 0.8744 | 0.9074 |
|  | M1 | 0.0513 | 0.0502 | 0.0441 | 0.0533 | 0.0383 | 0.0389 | 0.0558 | 0.0505 | 0.0446 | 0.0555 | 0.0254 | 0.0540 |
|  | M2 | 0.0000 | 0.0428 | 0.0133 | 0.0290 | 0.0055 | 0.0000 | 0.0000 | 0.0000 | 0.0427 | 0.0066 | 0.0424 | 0.0049 |
|  | M3 | 0.0053 | 0.0100 | 0.0059 | 0.0220 | 0.0179 | 0.0060 | 0.0143 | 0.0064 | 0.0125 | 0.0129 | 0.0285 | 0.0216 |
|  | M4 | 0.0002 | 0.0267 | 0.0045 | 0.0101 | 0.0068 | 0.0018 | 0.0023 | 0.0065 | 0.0000 | 0.0000 | 0.0292 | 0.0121 |
|  | M0 error | 0.0173 | 0.0512 | 0.0173 | 0.0176 | 0.0173 | 0.0173 | 0.0173 | 0.0173 | 0.0439 | 0.0260 | 0.0408 | 0.0173 |
|  | M1 error | 0.0173 | 0.0512 | 0.0173 | 0.0176 | 0.0173 | 0.0173 | 0.0173 | 0.0173 | 0.0439 | 0.0260 | 0.0408 | 0.0173 |
|  | M2 error | 0.0173 | 0.0512 | 0.0173 | 0.0176 | 0.0173 | 0.0173 | 0.0173 | 0.0173 | 0.0439 | 0.0260 | 0.0408 | 0.0173 |
|  | M3 error | 0.0173 | 0.0512 | 0.0173 | 0.0176 | 0.0173 | 0.0173 | 0.0173 | 0.0173 | 0.0439 | 0.0260 | 0.0408 | 0.0173 |
|  | M4 error | 0.0173 | 0.0512 | 0.0173 | 0.0176 | 0.0173 | 0.0173 | 0.0173 | 0.0173 | 0.0439 | 0.0260 | 0.0408 | 0.0173 |
| G1P | M0 | 0.8630 | 0.8418 | 0.8455 | 0.8490 | 0.8384 | 0.7878 | 0.8186 | 0.7312 | 0.7315 | 0.6975 | 0.6219 | 0.5804 |
|  | M1 | 0.1081 | 0.1259 | 0.1239 | 0.1175 | 0.1377 | 0.1679 | 0.1346 | 0.1436 | 0.1482 | 0.1340 | 0.1263 | 0.1239 |
|  | M2 | 0.0109 | 0.0153 | 0.0131 | 0.0183 | 0.0108 | 0.0265 | 0.0260 | 0.0525 | 0.0540 | 0.0640 | 0.0603 | 0.0684 |
|  | M3 | 0.0000 | 0.0048 | 0.0094 | 0.0012 | 0.0035 | 0.0071 | 0.0087 | 0.0384 | 0.0249 | 0.0380 | 0.0381 | 0.0630 |
|  | M4 | 0.0072 | 0.0061 | 0.0000 | 0.0004 | 0.0077 | 0.0016 | 0.0060 | 0.0125 | 0.0182 | 0.0326 | 0.0670 | 0.0541 |
|  | M5 | 0.0011 | 0.0004 | 0.0000 | 0.0022 | 0.0000 | 0.0050 | 0.0002 | 0.0147 | 0.0168 | 0.0171 | 0.0487 | 0.0565 |
|  | M6 | 0.0096 | 0.0057 | 0.0081 | 0.0114 | 0.0020 | 0.0040 | 0.0059 | 0.0070 | 0.0065 | 0.0169 | 0.0378 | 0.0537 |
|  | M0 error | 0.0547 | 0.0547 | 0.0547 | 0.0547 | 0.0547 | 0.0547 | 0.0547 | 0.0547 | 0.0547 | 0.0547 | 0.0596 | 0.0547 |
|  | M1 error | 0.0547 | 0.0547 | 0.0547 | 0.0547 | 0.0547 | 0.0547 | 0.0547 | 0.0547 | 0.0547 | 0.0547 | 0.0596 | 0.0547 |
|  | M2 error | 0.0547 | 0.0547 | 0.0547 | 0.0547 | 0.0547 | 0.0547 | 0.0547 | 0.0547 | 0.0547 | 0.0547 | 0.0596 | 0.0547 |
|  | M3 error | 0.0547 | 0.0547 | 0.0547 | 0.0547 | 0.0547 | 0.0547 | 0.0547 | 0.0547 | 0.0547 | 0.0547 | 0.0596 | 0.0547 |
|  | M4 error | 0.0547 | 0.0547 | 0.0547 | 0.0547 | 0.0547 | 0.0547 | 0.0547 | 0.0547 | 0.0547 | 0.0547 | 0.0596 | 0.0547 |
|  | M5 error | 0.0547 | 0.0547 | 0.0547 | 0.0547 | 0.0547 | 0.0547 | 0.0547 | 0.0547 | 0.0547 | 0.0547 | 0.0596 | 0.0547 |
|  | M6 error | 0.0547 | 0.0547 | 0.0547 | 0.0547 | 0.0547 | 0.0547 | 0.0547 | 0.0547 | 0.0547 | 0.0547 | 0.0596 | 0.0547 |
| GAP | M0 | 0.9538 | 0.8336 | 0.7730 | 0.7874 | 0.7652 | 0.7379 | 0.7156 | 0.6817 | 0.5672 | 0.5838 | 0.4031 | 0.4110 |
|  | M1 | 0.0439 | 0.1546 | 0.2076 | 0.1999 | 0.1984 | 0.2096 | 0.1982 | 0.1824 | 0.2184 | 0.1774 | 0.2042 | 0.2256 |
|  | M2 | 0.0000 | 0.0000 | 0.0000 | 0.0000 | 0.0000 | 0.0000 | 0.0000 | 0.0000 | 0.0000 | 0.0000 | 0.0000 | 0.0000 |
|  | M3 | 0.0023 | 0.0118 | 0.0193 | 0.0127 | 0.0364 | 0.0525 | 0.0862 | 0.1358 | 0.2145 | 0.2388 | 0.3927 | 0.3634 |
|  | M0 error | 0.0119 | 0.0405 | 0.0752 | 0.0228 | 0.0548 | 0.0615 | 0.0426 | 0.0463 | 0.0553 | 0.0931 | 0.0151 | 0.0557 |
|  | M1 error | 0.0119 | 0.0405 | 0.0752 | 0.0228 | 0.0548 | 0.0615 | 0.0426 | 0.0463 | 0.0553 | 0.0931 | 0.0151 | 0.0557 |
|  | M2 error | 0.0119 | 0.0405 | 0.0752 | 0.0228 | 0.0548 | 0.0615 | 0.0426 | 0.0463 | 0.0553 | 0.0931 | 0.0151 | 0.0557 |
|  | M3 error | 0.0119 | 0.0405 | 0.0752 | 0.0228 | 0.0548 | 0.0615 | 0.0426 | 0.0463 | 0.0553 | 0.0931 | 0.0151 | 0.0557 |
| PEP | M0 | 0.8619 | 0.8009 | 0.7842 | 0.7156 | 0.6269 | 0.6393 | 0.5644 | 0.5009 | 0.5024 | 0.5264 | 0.3996 | 0.3577 |
|  | M1 | 0.0682 | 0.1409 | 0.1436 | 0.1846 | 0.2394 | 0.2176 | 0.2210 | 0.2087 | 0.1692 | 0.1834 | 0.2225 | 0.2305 |
|  | M2 | 0.0568 | 0.0420 | 0.0497 | 0.0723 | 0.0960 | 0.0838 | 0.1214 | 0.1398 | 0.1493 | 0.1268 | 0.1735 | 0.1961 |
|  | M3 | 0.0131 | 0.0161 | 0.0225 | 0.0274 | 0.0376 | 0.0593 | 0.0933 | 0.1506 | 0.1791 | 0.1634 | 0.2044 | 0.2157 |
|  | M0 error | 0.0935 | 0.0935 | 0.0935 | 0.0935 | 0.0935 | 0.0935 | 0.0935 | 0.0935 | 0.0935 | 0.0935 | 0.0935 | 0.0935 |
|  | M1 error | 0.0935 | 0.0935 | 0.0935 | 0.0935 | 0.0935 | 0.0935 | 0.0935 | 0.0935 | 0.0935 | 0.0935 | 0.0935 | 0.0935 |
|  | M2 error | 0.0935 | 0.0935 | 0.0935 | 0.0935 | 0.0935 | 0.0935 | 0.0935 | 0.0935 | 0.0935 | 0.0935 | 0.0935 | 0.0935 |
|  | M3 error | 0.0935 | 0.0935 | 0.0935 | 0.0935 | 0.0935 | 0.0935 | 0.0935 | 0.0935 | 0.0935 | 0.0935 | 0.0935 | 0.0935 |
| R5P | M0 | 0.9025 | 0.7538 | 0.7432 | 0.6681 | 0.6768 | 0.5860 | 0.5479 | 0.4979 | 0.4305 | 0.4081 | 0.3600 | 0.3175 |
|  | M1 | 0.0663 | 0.1616 | 0.1745 | 0.2163 | 0.1800 | 0.2185 | 0.2072 | 0.2084 | 0.1717 | 0.1604 | 0.1304 | 0.1443 |
|  | M2 | 0.0201 | 0.0582 | 0.0519 | 0.0757 | 0.1032 | 0.1257 | 0.1434 | 0.1334 | 0.1794 | 0.1718 | 0.1855 | 0.2306 |
|  | M3 | 0.0013 | 0.0165 | 0.0129 | 0.0265 | 0.0293 | 0.0449 | 0.0724 | 0.1020 | 0.1249 | 0.1212 | 0.1409 | 0.1280 |
|  | M4 | 0.0083 | 0.0054 | 0.0161 | 0.0114 | 0.0077 | 0.0125 | 0.0213 | 0.0428 | 0.0492 | 0.0780 | 0.1127 | 0.1143 |
|  | M5 | 0.0014 | 0.0045 | 0.0014 | 0.0021 | 0.0031 | 0.0124 | 0.0079 | 0.0156 | 0.0442 | 0.0604 | 0.0705 | 0.0653 |
|  | M0 error | 0.0331 | 0.0331 | 0.0568 | 0.0331 | 0.0346 | 0.0363 | 0.0331 | 0.0535 | 0.0748 | 0.0331 | 0.0347 | 0.0640 |
|  | M1 error | 0.0331 | 0.0331 | 0.0568 | 0.0331 | 0.0346 | 0.0363 | 0.0331 | 0.0535 | 0.0748 | 0.0331 | 0.0347 | 0.0640 |
|  | M2 error | 0.0331 | 0.0331 | 0.0568 | 0.0331 | 0.0346 | 0.0363 | 0.0331 | 0.0535 | 0.0748 | 0.0331 | 0.0347 | 0.0640 |
|  | M3 error | 0.0331 | 0.0331 | 0.0568 | 0.0331 | 0.0346 | 0.0363 | 0.0331 | 0.0535 | 0.0748 | 0.0331 | 0.0347 | 0.0640 |
|  | M4 error | 0.0331 | 0.0331 | 0.0568 | 0.0331 | 0.0346 | 0.0363 | 0.0331 | 0.0535 | 0.0748 | 0.0331 | 0.0347 | 0.0640 |
|  | M5 error | 0.0331 | 0.0331 | 0.0568 | 0.0331 | 0.0346 | 0.0363 | 0.0331 | 0.0535 | 0.0748 | 0.0331 | 0.0347 | 0.0640 |

**Table S3: Metabolic Network and associated atom transitions:** The main reactions of cyanobacterial central metabolism and their atom transitions supplied to the INCA model

RUBP (abcde) + CO2 (f) -> 3PG (cde) + 3PG (fba)

3PG (abc) <-> GAP (abc)

GAP (abc) <-> DHAP (abc)

FBP (abcdef) <-> DHAP (cba) + GAP (def)

FBP (abcdef) <-> F6P (abcdef)

F6P (abcdef) <-> G6P (abcdef)

G6P (abcdef) -> RU5P (bcdef) + CO2 (a)

DHAP (abc) + E4P (defg) -> SBP (cbadefg)

SBP (abcdefg) -> S7P (abcdefg)

F6P (abcdef) <-> GAP (def) + EC3 (abc)

S7P (abcdefg) <-> E4P (defg) + EC3 (abc)

F6P (abcdef) <-> E4P (cdef) + EC2 (ab)

S7P (abcdefg) <-> R5P (cdefg) + EC2 (ab)

X5P (abcde) <-> GAP (cde) + EC2 (ab)

RU5P (abcde) <-> X5P (abcde)

RU5P (abcde) <-> R5P (abcde)

RU5P (abcde) -> RUBP (abcde)

RUBP (abcde) -> 3PG (cde) + 2PG (ba)

2PG (ab) -> GLYC (ab)

GLYC (ab) -> GOX (ab)

GOX (ab) + GOX (cd) -> GA (abd) + CO2 (c)

GA (abc) <-> 2PGA (abc)

3PG (abc) <-> 2PGA (abc)

2PGA (abc) <-> PEP (abc)

PEP (abc) -> PYR (abc)

PYR (abc) -> ACA (bc) + CO2 (a)

OAA (abcd) + ACA (ef) -> CIT (dcbfea)

CIT (abcdef) <-> ICI (abcdef)

ICI (abcdef) -> AKG (abcde) + CO2 (f)

AKG (abcde) -> SSA (bcde) + CO2 (a)

SSA (abcd) -> SUC (abcd)

SUC (abcd) <-> FUM (abcd)

FUM (abcd) <-> MAL (abcd)

MAL (abcd) <-> OAA (abcd)

PEP (abc) + CO2 (d) -> OAA (abcd)

MAL (abcd) -> PYR (abc) + CO2 (d)

0.4197*R5P + 3.579*ACA + 0.2171*E4P + 1.07*3PG + 0.5718*PEP + 2.728*PYR + 0.7506*OAA (abcd) + 0.7473*AKG + 0.21*GAP + 1.279*G1P -> 34.22*Biomass + 0.6313*FUM (abcd) + 1.536*dummy

G6P (abcdef) + dummy -> RU5P (bcdef) + CO2 (a)

GOX (ab) + GOX (cd) + dummy -> GA (abd) + CO2 (c)

PYR (abc) + dummy -> ACA (bc) + CO2 (a)

ICI (abcdef) + dummy <-> AKG (abcde) + CO2 (f)

AKG (abcde) + dummy -> SSA (bcde) + CO2 (a)

PEP (abc) + CO2 (d) -> OAA (abcd) + dummy

MAL (abcd) + dummy -> PYR (abc) + CO2 (d)

0*3PG (abc) -> 3PG.s (abc)

0*3PG.u (abc) -> 3PG.s (abc)

3PG.s -> sink

0*RU5P (abcde) -> RU5P.s (abcde)

0*RU5P.u (abcde) -> RU5P.s (abcde)

RU5P.s -> sink

0*PEP (abc) -> PEP.s (abc)

0*PEP.u (abc) -> PEP.s (abc)

PEP.s -> sink

0*GAP (abc) -> GAP.s (abc)

0*GAP.u (abc) -> GAP.s (abc)

GAP.s -> sink

G6P (abcdef) <-> G1P (abcdef)

0*DHAP (abc) -> DHAP.s (abc)

0*DHAP.u (abc) -> DHAP.s (abc)

DHAP.s -> sink

CO2.l (a) -> CO2.x (a)

CO2.u (a) -> CO2.x (a)

*GAP (def) + PYR (abc) -> DOXP (defba) + CO2 (c)

*DOXP (abcde) + DOXP (fghij) -> LIM (abcdefghij)

CO2.x (a) -> CO2 (a)

CO2.u (a) + dummy -> CO2 (a)

*Applies only to the LS strain

**Table S4: Simulated flux values and bounds.** Parameter continuation was used to determine the upper and lower bound of each reaction flux to a 95% confidence interval. Biomass: 0.4197*R5P + 3.579*ACA + 0.2171*E4P + 1.07*3PG + 0.5718*PEP + 2.728*PYR + 0.7506*OAA + 0.7473*AKG + 0.21*GAP + 1.279*G1P -> 34.22*Biomass + 0.6313*FUM + 1.536*dummy

| Reaction | WT Flux | WT StdErr | WT LB | WT UB | LS Flux | LS StdErr | LS LB | LS UB |
| --- | --- | --- | --- | --- | --- | --- | --- | --- |
| RUBP + CO2 -> 3PG + 3PG | 72.68 | 3.71 | 63.99 | 82.30 | 70.96 | 3.65 | 63.08 | 78.96 |
| 3PG <-> GAP | 127.97 | 6.53 | 112.80 | 143.73 | 124.96 | 6.43 | 111.03 | 139.18 |
| GAP <-> DHAP | 51.42 | 2.63 | 45.26 | 57.89 | 50.19 | 2.59 | 44.63 | 56.02 |
| FBP <-> DHAP + GAP | -27.07 | 1.38 | -30.53 | -23.82 | -26.42 | 1.36 | -29.42 | -23.48 |
| FBP <-> F6P | 27.07 | 1.38 | 23.82 | 30.53 | 26.42 | 1.36 | 23.48 | 29.42 |
| F6P <-> G6P | 2.33 | 0.12 | 2.05 | 2.78 | 2.26 | 0.12 | 2.01 | 2.53 |
| G6P -> RU5P + CO2 | 0.00 | 0.00 | 0.00 | 0.38 | 0.00 | 0.00 | 0.00 | 0.16 |
| DHAP + E4P -> SBP | 24.35 | 1.24 | 21.31 | 27.51 | 23.77 | 1.22 | 21.14 | 26.48 |
| SBP -> S7P | 24.35 | 1.24 | 21.31 | 27.51 | 23.77 | 1.22 | 21.14 | 26.48 |
| F6P <-> GAP + EC3 | 0.00 | 0.00 | 0.00 | 0.61 | 0.00 | 0.00 | 0.00 | 0.51 |
| S7P <-> E4P + EC3 | 0.00 | 0.00 | -0.61 | 0.00 | 0.00 | 0.00 | -0.51 | 0.00 |
| F6P <-> E4P + EC2 | 24.74 | 1.26 | 21.80 | 27.93 | 24.16 | 1.24 | 21.47 | 26.90 |
| S7P <-> R5P + EC2 | 24.35 | 1.24 | 21.46 | 27.26 | 23.77 | 1.22 | 21.12 | 26.49 |
| X5P <-> GAP + EC2 | -49.09 | 2.51 | -55.51 | -43.32 | -47.93 | 2.47 | -53.33 | -42.60 |
| RU5P <-> X5P | -49.09 | 2.51 | -55.51 | -43.32 | -47.93 | 2.47 | -53.33 | -42.60 |
| RU5P <-> R5P | -23.59 | 1.20 | -26.69 | -20.79 | -23.03 | 1.19 | -25.64 | -20.47 |
| RU5P -> RUBP | 72.68 | 3.71 | 64.11 | 81.98 | 70.96 | 3.65 | 63.08 | 78.99 |
| RUBP -> 3PG + 2PG | 0.00 | 0.00 | 0.00 | 0.46 | 0.00 | 0.00 | 0.00 | 0.83 |
| 2PG -> GLYC | 0.00 | 0.00 | 0.00 | 0.46 | 0.00 | 0.00 | 0.00 | 0.83 |
| GLYC -> GOX | 0.00 | 0.00 | 0.00 | 0.46 | 0.00 | 0.00 | 0.00 | 0.83 |
| GOX + GOX -> GA + CO2 | 0.00 | 0.00 | 0.00 | 0.23 | 0.00 | 0.00 | 0.00 | 0.41 |
| GA <-> 2PGA | 0.00 | 0.00 | 0.00 | 0.23 | 0.00 | 0.00 | 0.00 | 0.41 |
| 3PG <-> 2PGA | 15.45 | 0.79 | 13.51 | 17.46 | 15.07 | 0.78 | 13.37 | 16.74 |
| 2PGA <-> PEP | 15.45 | 0.79 | 13.58 | 17.32 | 15.07 | 0.78 | 13.39 | 16.78 |
| PEP -> PYR | 9.86 | 2.04 | 7.95 | 11.68 | 0.00 | 0.00 | NaN | 12.15 |
| PYR -> ACA + CO2 | 7.87 | 0.40 | 6.92 | 8.82 | 7.66 | 0.40 | 6.80 | 8.52 |
| OAA + ACA -> CIT | 1.36 | 0.07 | 1.19 | 1.59 | 1.32 | 0.07 | 1.17 | 1.49 |
| CIT <-> ICI | 1.36 | 0.07 | 1.19 | 1.59 | 1.32 | 0.07 | 1.17 | 1.49 |
| ICI -> AKG + CO2 | 1.36 | 0.07 | 1.19 | 1.59 | 1.32 | 0.07 | 1.17 | 1.47 |
| AKG -> SSA + CO2 | 0.00 | 0.00 | 0.00 | 0.13 | 0.00 | 0.00 | 0.00 | 0.09 |
| SSA -> SUC | 0.00 | 0.00 | 0.00 | 0.13 | 0.00 | 0.00 | 0.00 | 0.09 |
| SUC <-> FUM | 0.00 | 0.00 | 0.00 | 0.13 | 0.00 | 0.00 | 0.00 | 0.11 |
| FUM <-> MAL | 1.15 | 0.06 | 1.01 | 1.32 | 1.12 | 0.06 | 0.99 | 1.27 |
| MAL <-> OAA | -1.82 | 1.97 | -3.46 | -0.53 | -11.41 | 0.59 | -12.67 | 0.05 |
| PEP + CO2 -> OAA | 4.55 | 1.98 | 3.20 | 6.42 | 14.06 | 0.72 | 3.57 | 15.66 |
| MAL -> PYR + CO2 | 2.97 | 1.97 | 1.64 | 4.94 | 12.52 | 0.65 | 1.67 | 13.95 |
| Biomass | 1.82 | 0.09 | 1.60 | 2.06 | 1.77 | 0.09 | 1.57 | 1.97 |
| 0*3PG -> 3PG.s | 0.86 | 0.04 | 0.78 | 0.96 | 0.77 | 0.06 | 0.64 | 0.89 |
| 0*3PG.u -> 3PG.s | 0.14 | 0.04 | 0.04 | 0.22 | 0.23 | 0.06 | 0.11 | 0.36 |
| 3PG.s -> sink | 1.00 | 0.00 | 1.00 | 1.00 | 1.00 | 0.00 | 1.00 | 1.00 |
| 0*RU5P -> RU5P.s | 0.68 | 0.01 | 0.66 | 0.71 | 0.58 | 0.02 | 0.54 | 0.62 |
| 0*RU5P.u -> RU5P.s | 0.32 | 0.01 | 0.29 | 0.34 | 0.42 | 0.02 | 0.38 | 0.46 |
| RU5P.s -> sink | 1.00 | 0.00 | 1.00 | 1.00 | 1.00 | 0.00 | 1.00 | 1.00 |
| 0*PEP -> PEP.s | 0.86 | 0.02 | 0.81 | 0.90 | 0.60 | 0.04 | 0.51 | 0.68 |
| 0*PEP.u -> PEP.s | 0.15 | 0.02 | 0.10 | 0.19 | 0.40 | 0.04 | 0.32 | 0.49 |
| PEP.s -> sink | 1.00 | 0.00 | 1.00 | 1.00 | 1.00 | 0.00 | 1.00 | 1.00 |
| 0*GAP -> GAP.s | 0.81 | 0.01 | 0.80 | 0.83 | 0.70 | 0.01 | 0.67 | 0.72 |
| 0*GAP.u -> GAP.s | 0.19 | 0.01 | 0.17 | 0.20 | 0.30 | 0.01 | 0.28 | 0.33 |
| GAP.s -> sink | 1.00 | 0.00 | 1.00 | 1.00 | 1.00 | 0.00 | 1.00 | 1.00 |
| G6P <-> G1P | 2.33 | 0.12 | 2.05 | 2.63 | 2.26 | 0.12 | 2.01 | 2.52 |
| 0*DHAP -> DHAP.s | 0.81 | 0.01 | 0.80 | 0.82 | 0.66 | 0.02 | 0.62 | 0.69 |
| 0*DHAP.u -> DHAP.s | 0.19 | 0.01 | 0.18 | 0.20 | 0.34 | 0.02 | 0.31 | 0.38 |
| DHAP.s -> sink | 1.00 | 0.00 | 1.00 | 1.00 | 1.00 | 0.00 | 1.00 | 1.00 |
| CO2.l -> CO2.x | 62.23 | 3.18 | 54.61 | 69.67 | 60.76 | 3.13 | 54.03 | 67.53 |
| CO2.u -> CO2.x | 0.00 | 0.00 | 0.00 | 0.34 | 0.00 | 0.00 | 0.00 | 0.32 |
| GAP + PYR -> DOXP + CO2 | N/A | N/A | N/A | N/A | 0.04 | 0.00 | 0.04 | 0.04 |
| DOXP + DOXP -> LIM | N/A | N/A | N/A | N/A | 0.02 | 0.00 | 0.02 | 0.02 |
| CO2.x -> CO2 | 62.23 | 3.18 | 54.87 | 70.38 | 60.76 | 3.13 | 53.94 | 67.52 |
| CO2.u + dummy -> CO2 | 2.79 | 0.14 | 2.46 | 3.16 | 2.72 | 0.14 | 2.41 | 3.02 |

**Figure S2: Growth curve of Synechococcus 7002 WT strain and LS strain.** The labeling and quenching experiment was performed at an OD_730_ of around 0.8. We converted the average growth rate between the data points indicated by the arrow to a biomass accumulation rate in terms of umol L^-1^ hr^-1^ using a conversion factor derived by [1].

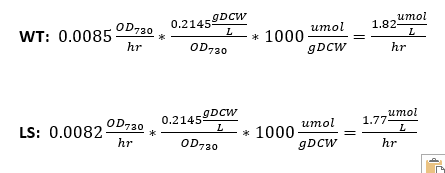

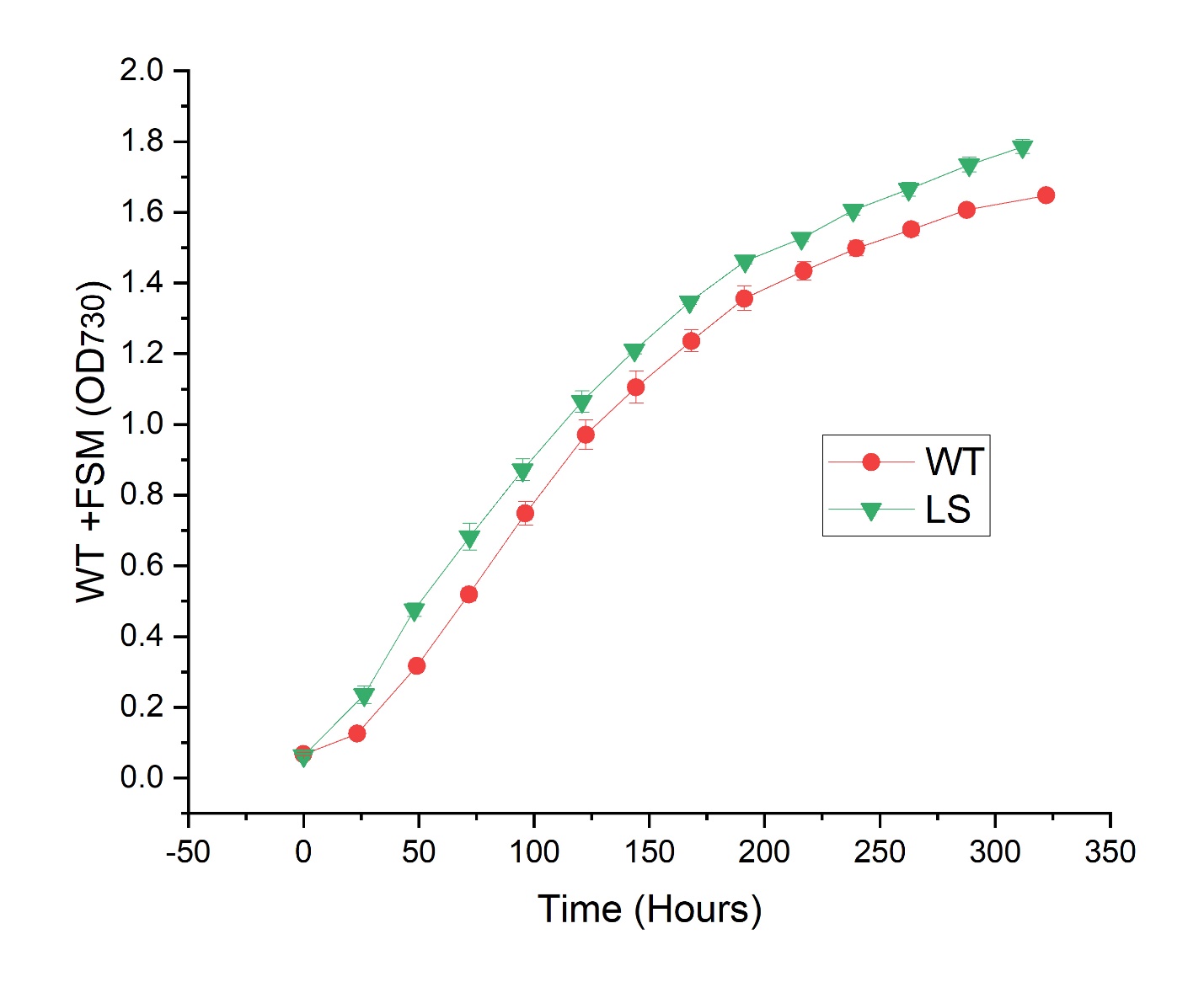

1.	Selão, T.T., et al., *Growth and selection of the cyanobacterium Synechococcus sp. PCC 7002 using alternative nitrogen and phosphorus sources.* Metabolic Engineering, 2019. **54**: p. 255-263.
